# Sequence-specific dynamics of DNA response elements and their flanking sites regulate the recognition by AP-1 transcription factors

**DOI:** 10.1101/2020.05.31.125989

**Authors:** Johanna Hörberg, Kevin Moreau, Anna Reymer

**Affiliations:** Department of Chemistry and Molecular Biology, University of Gothenburg, Gothenburg 40530, Sweden

## Abstract

Activator proteins 1 (AP-1) comprise one of the largest families of eukaryotic basic leucine zipper transcription factors. Despite advances in the characterization of AP-1 DNA-binding sites, our ability to predict new binding sites and explain how the proteins achieve different gene expression levels remains limited. Here we address the role of sequence-specific DNA dynamics for stability and specific binding of AP-1 factors, using microseconds long molecular dynamics simulations. As a model system, we employ yeast AP-1 factor Yap1 binding to three different response elements from two genetic environments. Our data show that Yap1 actively exploits the sequence-specific plasticity of DNA within the response element to form stable protein-DNA complexes. The stability also depends on the four to six flanking nucleotides, adjacent to the response elements. The flanking sequences modulate the conformational adaptability of the response element, making it more shape-efficient to form specific contacts with the protein. Bioinformatics analysis of differential expression of the studied genes supports our conclusions: the stability of Yap1-DNA complexes, modulated by the flanking environment, influences the gene expression levels. Our results provide new insights into mechanisms of protein-DNA recognition and the biological regulation of gene expression levels in eukaryotes.

## Introduction

Activator proteins 1 (AP-1) comprise one of the largest and most evolutionary conserved families of transcription factor proteins (TFs) in eukaryotes, which regulate among other cellular stress responses, cell differentiation and cell proliferation.^1,2^ The AP-1 factors constitute a subgroup of basic leucine zippers (BZIPs),^3^ which control gene transcription by binding as homo- or heterodimers^2,4,5^ to specific DNA targets known as AP-1 response elements (ARE). The AP-1 proteins achieve their DNA selectivity predominantly through a direct readout mechanism, when a highly conserved five-residues-motif of the protein basic region, **N**XX**AA**XX**CR**, recognizes the ARE-DNA half-site.^3^ Despite the high sequence homology of the basic regions, AP-1 factors recognize diverse pseudo-palindromic and palindromic AREs, as well as emergent sites containing only a consensus-like half-site.^3,5–7^ The lengths of AP-1 response elements also vary. Commonly, AP-1 proteins recognize DNA targets of seven or eight base pairs (b.p.), where the experimentally derived consensus sequence corresponds to a seven b.p. pseudo-palindrome, TGACTCA.^3^ However, several members of the AP-1 family show preference for longer AREs of 13-14 b.p.^6,8,9^ This diversity of ARE-sequences hampers our understanding of how AP-1 factors achieve their DNA binding selectivity, and consequently the transcription regulation specificity.

The binding of AP-1 factors induce no major DNA deformation,^3,10^ as observed in experimentally solved structures of AP-1 protein-DNA complexes, however the proteins can potentially exploit local sequence-specific structural variations of DNA.^11^ The early analysis of crystallographic structures of protein-DNA complexes, by Olsson and colleagues,^12^ proposed that the most polymorphic pyrimidine(Y)-purine(R) dinucleotide steps could act as flexible “hinges” facilitating the conformational adjustment of DNA to its protein-partner. However, due to the limited number of protein-DNA structures, the study focused on the nearest-neighbour effects. Progress in atomistic molecular dynamics (MD) and in the force fields for nucleic acids, provided further insights into the role of DNA sequence-specific structural dynamics for protein recognition.^13^ Microsecond-long MD simulations of DNA oligomers by ABC consortium,^14,15^ confirmed that certain dinucleotides (YR and RR) oscillate between conformational substates. The studies reported: the local DNA polymorphism is highly heterogeneous and sequence-specific, and is coupled to the tetranucleotide^14^ or even hexanucleotide level,^15^ depending both on the nucleotide composition of the central dinucleotide step and its flanking environment. As a result, once bound by a protein the local plasticity of DNA can finalize the DNA transition into a bioactive conformation, as reported by an extensive MD study by Orozco and colleagues.^16^

Another important, but much less explored aspect that can modulate the specific binding of AP-1 factors is flanking sequences outside the response elements.^17^ The high-throughput study by Steger and colleagues^18^ focusing on another subfamily of BZIPs, CREBP transcription factors, showed a strong preference for 5’-R and 3’-Y flanking nucleotides, directly adjacent to the response elements. Bansal and colleagues^19^ further highlighted the importance of the flanking regions for several transcription factors, including the BZIP proteins Fos-Jun (which prefer 5’-RR and 3’-YY-flanks) and NFIL3 (which prefer YR for both 5’- and 3’-flanks). Both studies proposed that DNA shape readout could contribute to BZIP-proteins binding specificity, since only nonspecific contacts with the DNA backbone of the flanking nucleotides were found in the crystal structures of the BZIP-DNA complexes. These studies provide evidence that the flanking sequences could play a regulatory role in transcription factors specific binding to DNA, but the underlying molecular mechanism and the number of contributing flanking nucleotides remains to be discovered.

In this paper, we explore mechanistic aspects of the AP-1-DNA recognition process, focusing on the local DNA sequence-specific dynamics, variations in the response elements and flanking environments, using atomistic molecular dynamics simulation in the microsecond range. As a model system, we employ Yap1 transcription factor,^7^ a member of Saccharomyces cerevisiae AP-1 family, which regulates genes involved in oxidative stress responses and cell detoxification. Differently from AP-1 transcription factors in other eukaryotic organisms, the basic region of Yap1 contains glutamine and phenylalanine instead of alanine and cysteine (**N**xxAQxx**F**R).^3^ The phenylalanine presence suggests an increased preference for 5’-TT dinucleotide step at the extremities of Yap1 response element.^3,7^ However, the response elements identified in promoters of Yap1-modulated genes remain diverse:^7,20–24^ TTACTAA, TTACGTAA, TGA(C/G)TAA, T(T/G)ACAAA, etc. For our computational study, we select three different Yap1 response elements from promoter regions of two genes, to account for a variation in the flanking sequences. We observe that the local variations of the helical parameters, shift and slide, within the DNA response element modulate the binding specificity of Yap1, and are sequence-specific. Our data also show that four to six nucleotides, flanking the response elements, influence how easily shift and, to a lesser extent, slide can adjust within the response element to achieve the bioactive conformation upon the protein binding. This sequence-specific plasticity, regulated by the flanking environment, we propose, play a unique role in the recognition process as well as in the stability of the protein-DNA complexes, modulating the firing strength of the corresponding promoters. Our mechanistic findings correlate well with the expression levels of the two selected genes, and of the model-reporter genes containing one instance of the selected Yap1 response elements.

## Methods

### Systems Studied

We study 12 systems: Yap1 bound to three different response elements (YRE1: TTACTAA, YRE2: TTACGTAA, YRE3: TGACAAA) in two genomic environments (ATR1, OYE2),^25^ as well as unbound DNA in B-form for each corresponding system. The native DNA sequences are obtained from the NCBI database^26^ (see Figure S1 for their position relative to the transcription starting site) with the Gene ID 854924 and 856584 for ATR1 and OYE2, respectively. The six studied 23-mer DNA sequences are listed in Table 1.

**Table 1:**
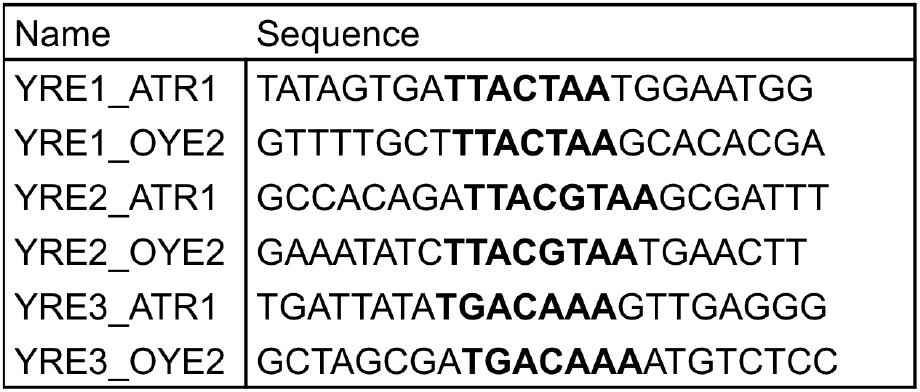
The six studied 23-mer DNA sequences in two genomic environments (ATR1 and OYE2) containing the response elements in bold

### Homology Modelling

The Yap1 homodimer is derived through homology modelling in YASARA.^27^ The Fasta file for the Yap1 BZIP domain (residues 63-130, obtained from Uniprot^28^ with ID P19880, is subjected to the comparative modelling tool in YASARA, using the high resolution crystal structure of Pap1 (PDB ID: 1GD2),^3^ the closest orthologue of Yap1, as a template. Quality assessment of the Yap1 homology model, performed by YASARA, where the derived model obtained an optimal score, indicates a high-resolution structure. For the complete parameter sets used for the homology modelling see Table S1.

### Protein-DNA Docking

To derive the six Yap1-DNA complexes, protein-DNA docking is performed using HDOCK webserver.^29^ HDOCK performs rigid macromolecular docking, using a hybrid algorithm of template-based modelling and *ab initio* free docking. For the docking, Yap1 is defined as the receptor and B-DNA, containing the response elements (TTACTAA, TTACGTAA, TGACAAA) in the native flanking environment (ATR1, OYE2), is defined as the ligand. Using defaults settings positioned Yap1 to enable interactions with the response elements among the top-10 scored complexes. The derived Yap1-DNA complexes and free DNA in B-form are subsequently pre-processed using the modelling program JUMNA.^30^

### Molecular Dynamics Simulations

All molecular dynamics (MD) simulations are performed using the MD engine GROMACS v2018.1.^31^ For each simulation a combination of AMBER 14SB^32^ and Parmbc1^33^ force fields is used for the protein and DNA, respectively. The Yap1-DNA complexes and free DNA oligomers are separately solvated in triclinic rectangular periodic boxes by SPC/E water molecules^34^ with a buffer distance of 15 Å to the walls. Each system is neutralized by K+ counterions. Additional K+ and Cl-ions are then added to reach a physiological salt-concentration of 150 mM. Applying periodic boundary conditions, each system is subjected to energy minimization with 5000 steps of steepest descent, followed by 500 ps equilibration-runs with week position restraints on heavy solute atoms (1000 kJ/mol) in the NVT and NPT ensembles, adjusting temperature and pressure to 300 K and 1 atm. Releasing the restraints, 1.1 microsecond simulations are then carried out at constant pressure and temperature (1 atm and 300 K). Temperature is controlled by a weak-coupling thermostat^35^ with a coupling constant of 0.2 ps and pressure is controlled by an isotropic Parrinello-Rahman barostat^36^ with a coupling constant of 2 ps. All bonds involving hydrogen atoms are constrained with the LINCS algorithm,^37^ allowing the time step of 2 fs. Electrostatic interactions are treated with the Particle Mesh Ewald summation method^38^ using a short-range cutoff of 10 Å. The van-der-Waals forces are also truncated at 10 Å with added long-range corrections. The neighbour pair list for nonbonded interactions is updated every 20th step through the Verlet cutoff scheme.^39^ Centre of mass movement is removed every 0.2 ps to eliminate translational kinetic energy build-up.^40^

### Conformational Analysis

The generated trajectories are processed using CPPTRAJ program^41^ from AMBERTOOLS 16 software package. The first 100 ns of each trajectory are discarded as equilibration. Subsequently, Curves+, Canal, and Canion programs^42^ are used to derive the helical parameters, backbone torsional angles, groove geometry parameters and ion distributions for each trajectory snapshot extracted every ps.

### Contact-Network Analysis

Analysis of the protein-DNA contacts network for the six Yap1-DNA complexes is performed using CPPTRAJ^41^ for each trajectory snapshot extracted at 1 ps intervals. Protein-DNA contacts present for less than 10% of the trajectories are excluded. The protein-DNA contacts are characterized by pairs of residues, divided into ‘specific’, i.e. interactions formed between the protein side chains and DNA bases, and ‘non-specific’, i.e. interactions formed with at least one of the molecules’ backbones. The contacts formed between each protein-DNA residue pair are summed, where for simplicity, the contribution of each contact is set to 1. The distance limit of a hydrogen bond interaction is ≤ 4 Å between the relevant heavy atoms, and the angle limit is ≥ 135° at the intervening hydrogen atom. For a salt bridge, the limit is ≤ 4.0 Å between the end-group nitrogen of lysine and arginine, and the DNA phosphate group. For a hydrophobic contact, the limit is ≤ 6Å between the centre of mass of hydrophobic residues (Ala, Ile, Leu, Met, Phe, Trp, and Cys) and DNA bases. The derived time series of Yap1-DNA interactions allow construction of dynamic contacts maps for specific and non-specific contacts, characterizing the binding specificity and the stability of the Yap1-DNA complexes along the trajectories.

### BZIP-DNA Helical Parameters Motifs

Crystal structures of BZIP-DNA complexes bound to different response elements (TGACTCA, TGACGTCA, TTACGTAA) are downloaded from the Protein Data Bank,^43^ and analysed for helical parameters for the BZIP-DNA complexes and their corresponding naked DNA in B-form by Curves+.^42^

### Differential expression data mining

The differential expression levels of the two Yap1 target genes, ATR1 and OYE2, across different time-points are derived from the study by Salin *et al*.^44^ The data are downloaded from the ArrayExpress website through the access number E-TABM-439. The expression level profiles for the two genes are derived by calculating the log2 ratio between WT and the *yap1Δ* mutant under the selenite stress.

### YRE analysis on random promoters’ study

Data for randomly generated promoters with associated expression levels are derived from the study by Boer *et a*l.^45^, obtained from NCBI’s gene expression omnibus under the access number GSE104878. Internally developed R-scripts are used to isolate promoters possessing only one instance of YRE decorated with at least four of the adjacent 5’- and 3’-flanking nucleotides from the ATR1 or OYE2-environments (**NNNN**YRE**NNNN**), and to analyse the associated expression data.

### Additional Information

MatLab software is used for the post-processing and plotting of all data. USCF Chimera^46^ is used for creating the molecular graphics.

## Results and discussion

### Yap1-DNA contacts

To address the molecular mechanism of DNA recognition by AP-1 transcription factors, we study Yap1 transcription factor binding to three different Yap1 response elements (YRE), YRE1: TTACTAA, YRE2: TTACGTAA, and YRE3: TGACAAA. To identify the role of the flanking environment outside the response elements, we extract DNA sequences from two Yap1-regulated genes, ATR1 and OYE2, which contain the three YREs (Table 1, Figure S1). To design the 3-D structure of DNA-binding BZIP-domain of Yap1, we use the homology modelling approach with the high-resolution crystallographic structure of Pap1 (PDB ID: 1GD2)^3^ the closest orthologue of Yap1 as the template. To derive the structures of Yap1-DNA complexes we perform macromolecular docking with HDOCK webserver,^29^ which uses a hybrid algorithm of template-based modelling and *ab initio* free docking. And finally, to generate the dynamic profiles of macromolecular interactions and assess complexes stability, we subject each Yap1-YRE systems and the corresponding naked DNA oligonucleotides to a microsecond long all-atom MD simulation.

Our simulations show that, in analogy to other AP-1 transcription factors, Yap1 utilizes the five-residues-motif (**N**xxAQxx**F**R):^3^ Asn74, Ala77, Gln78, Phe81 and Arg82 to form the base-specific contacts with the three DNA response elements (Figure 1). We observe both similarities and differences in the specific contacts formed by Yap1, depending on the YRE and its flanking sites. The similarities include the contacts by Phe81, which recognizes the outer 5’-TT/TG step; by Arg82, which favours the central (C)G b.p./b.p. step; and by Ala77, which interacts with the first thymine of the outer 5’-TT/TG step. The differences include the contacts formed by Asn74 and Gln78 that exhibit a considerable variation in the specific contacts within the three YREs, which are also monomer-specific. For instance, Gln78 forms a number of hydrogen bonds with the TA/GA step (T**TA**CTAA, T**TA**CGTAA, and T**GA**CAAA) and its complementary TA/CT step on the opposite strand. Asn74 forms either monodentate/bidentate contacts with the outer 3’-AA step (TTACT**AA**, TTACGT**AA**, and TGACA**AA**), or cross-bridging contacts involving cytosine of the 3’-CA step and thymine of the 5’-TG step on the opposite strand (**T**GACAAA/TTTGT**C**A). The scope of the hydrogen bond contacts exploited by Asn74 and Gln78 suggests a compelling molecular mechanism explaining how Yap1 can recognize a variety of YREs. For further details of the intermolecular contacts, see Figure 1A-B.

**Figure 1.**
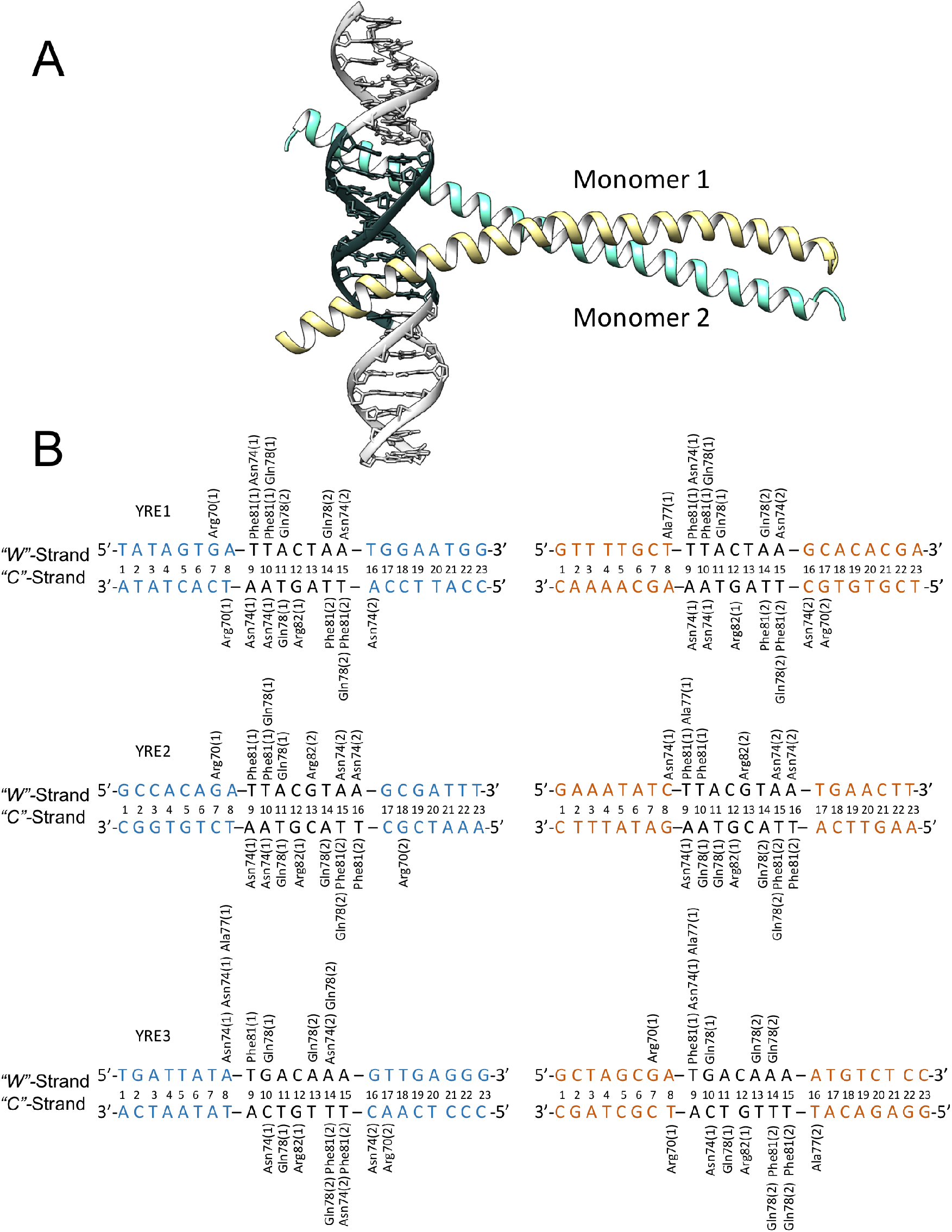
**A.** Homology model of Yap1 (monomer1 in yellow, monomer 2 in turquoise) bound to DNA, where YRE is highlighted with dark grey. **B.** Specific contacts between Yap1 and the three YREs (YRE1: TTACTAA, YRE2: TTACGTAA, YRE3: TGACAAA) in two genomic environments (ATR1: blue, OYE2: orange) observed in one microsecond long MD simulations.

### Impact of DNA Flanking Environment

In addition to the composition of the response element, we observe a significant impact of the flanking environment on the strength and specificity of Yap1-DNA contacts (Figure 2A-C). Since the Yap1-YRE recognition is dominated by the direct readout mechanism, we hypothesize that the number of specific protein-DNA contacts, namely the contacts formed between the protein side chains and DNA bases, represents the molecular selectivity. Whereas the number of the total protein-DNA contacts represents the complex stability. We characterize the strength of the protein-DNA interactions by pairs of residues, where for each pair we sum up all hydrogen bonds, salt bridges, and hydrophobic (apolar) interactions. For simplicity, the contribution of a single bond of each type is set to 1, since the energy cost varies depending on the nature of interacting atoms, the bond geometry, and the surrounding environment.

**Figure 2.**
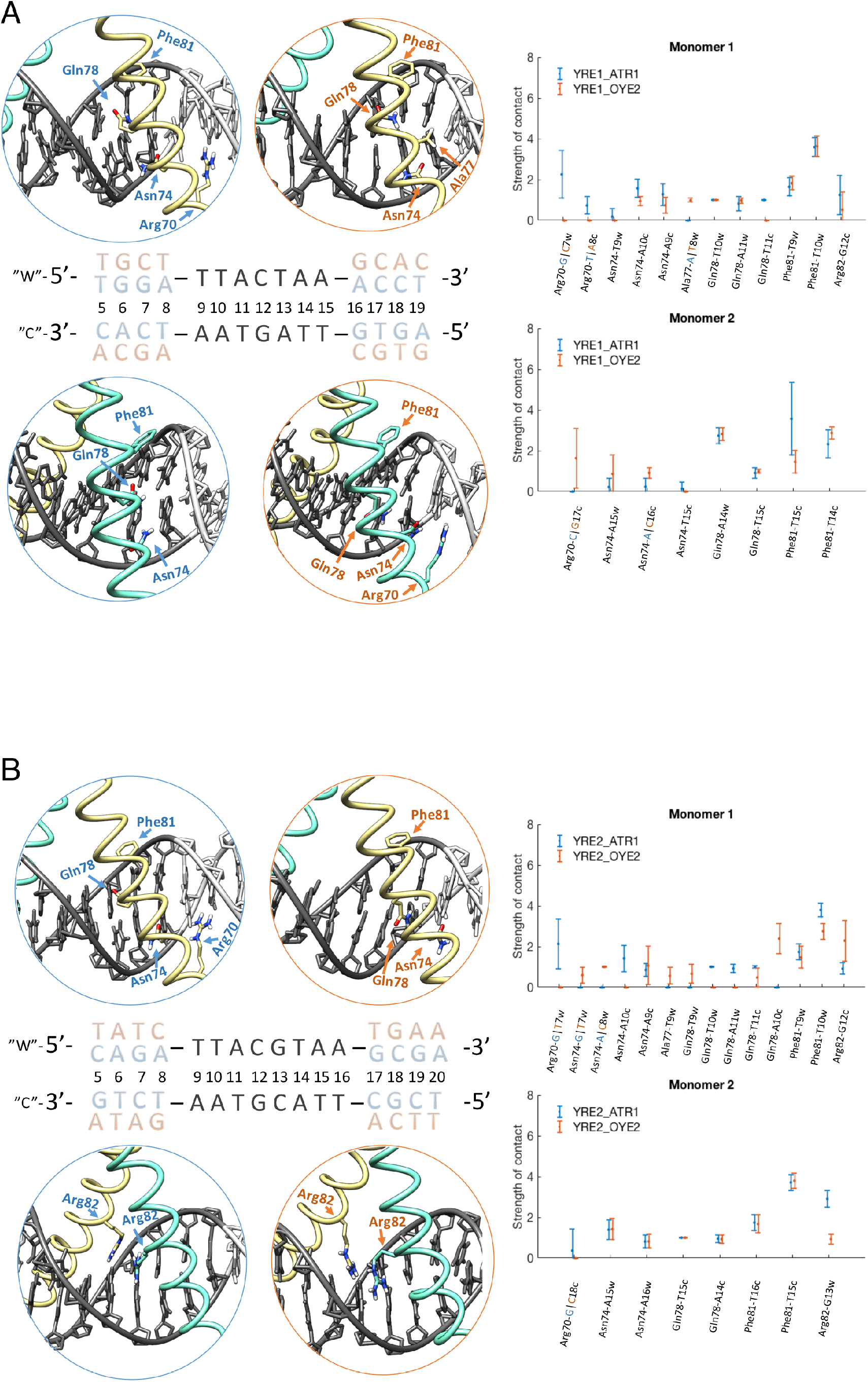
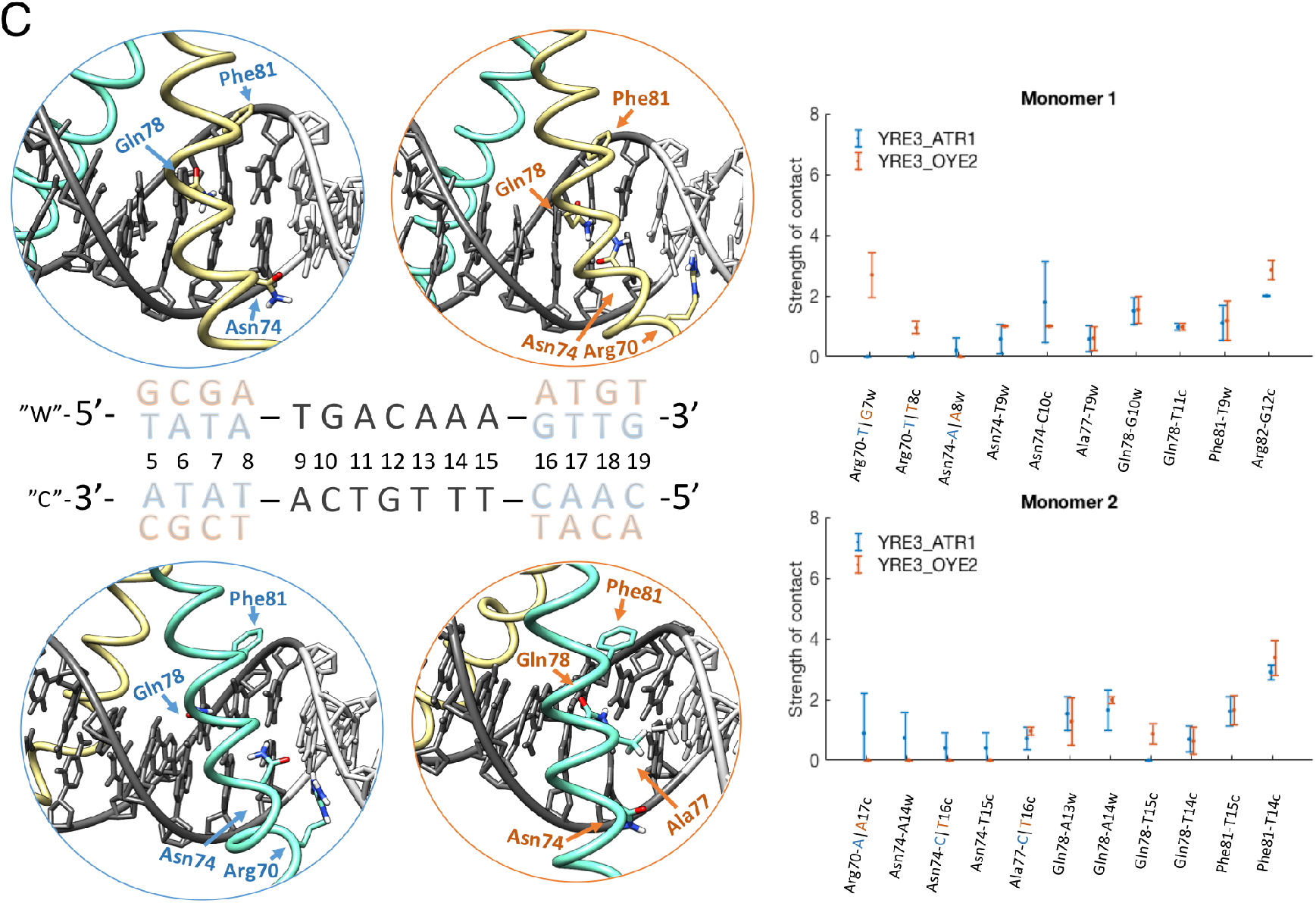
Specific contacts between Yap1 five-residues recognition motif (**R**xxx**N**xx**AQ**xx**FR**) and the three YREs in two genomic environments, ATR1 and OYE2. For the Yap1-DNA complexes, Yap1 monomer 1 is in yellow, Yap1 monomer 2 is in turquoise, and DNA is in grey, where YRE is highlighted with dark-grey. The ATR1-environment is blue-marked and the OYE2-environment is orange-marked. The plots show the strength of specific contacts exploited by Yap1 monomers in two genomic environments. We define contact strength by pairs of residues, i.e. for each Protein-DNA residue pair we sum all the contacts involving the protein residue side chains and the DNA bases. **A.** Yap1-YRE1. **B.** Yap1-YRE2. **C.** Yap1-YRE3.

We observe that for pseudo-palindromic YRE1 (TTACTAA), in two genomic environments ATR1 and OYE2 (Figure 2A, Figure S2), Yap1 monomer 1 exhibits similar strength of contacts formed by Gln78 and Phe81. The ATR1-environment contributes, however, to stronger Asn74-YRE1 and Arg82-YRE1 contacts. Moreover, in the ATR1-environment, YRE1 is surrounded by the adjacent 5’-GA flanking step (**GA**-TTACTAA) that creates an additional specific contact with Arg70 (**R**xxxNxxAQxxFR), a semi-conserved residue among the BZIP-families. The Arg70-G contacts allow for a tighter association of monomer 1 with the YRE1 half-site, stabilizing the specific contacts exploited by the five-residues-motif (**N**xx**AQ**xx**FR**). In contrast, in the OYE2-environment, YRE1 is surrounded by the adjacent 5’-CT step (**CT**-TTACTAA), where the flanking thymine, although contributing to additional hydrophobic contacts with Ala77 (Nxx**A**QxxFR), sterically hinders the tight association of monomer 1 with the YRE1 half-site (Figure 2A).

For Yap1 monomer 2, we observe noticeable differences in the specific contacts between the two genomic environments (Figure 2A). For the “Crick”-strand (3’->5’ DNA direction) in the OYE2-environment, YRE1 is flanked by the 5’-GC step, which results in Arg70-G and Asn74-C specific interactions (**GC-**TTAGTAA). This in turn, stabilizes the binding orientation of monomer 2, providing similar strengths of the Phe81-T interactions as for monomer 1. In contrast, in the ATR1-environment, YRE1 is flanked by the 5’-CA step (**CA**-TTAGTAA), which results in a less stable contact with Asn74. Thus, we observe higher fluctuations in the strength of specific contacts exploited by monomer 2 that, in particular, alter the Phe81-T interactions. Monomer 2 in the ATR1-environment does not follow the trend of Phe81 interacting with the outer 5’-TT step of YRE1 (**TT**AGTAA), where the CB atom forms hydrophobic contacts with the first thymine, and the aromatic ring forms contacts with the second thymine. Instead, sliding of monomer 2 positions the aromatic ring of Phe81 in-between the two thymines, with a preference towards the first thymine (Figure 2A, Figure S2). There are also some similarities: both genomic environments result in similar Gln78-YRE1 contacts, however, not symmetric with respect to the Gln78 contacts of monomer 1. Instead of interacting with the TA step (TTAC**TA**A), Gln78 of monomer 2 forms contacts with the outer 3’-AA step (TTACT**AA**). Consequently, this rearrangement of Gln78 pushes Asn74 towards interactions with the flanking environment (Figure 2A).

In the case of palindromic YRE2 (TTACGTAA), we observe greater differences in specific contacts exploited by Yap1 monomer 1 than by monomer 2 (Figure 2B, Figure S3). In the ATR1-environment, YRE2 is similarly to YRE1 flanked by 5’-GA step, which provides additional Arg70-G contacts and positions monomer 1 deeper into the major groove of YRE2 half-site. The specific contacts between the five-residues-motif (**N**xxx**AQ**xx**FR**) of monomer 1 and the YRE2 half-site follow the trends described for YRE1 in the same genomic environment. In contrast, in the OYE2-environment, YRE2 is surrounded by the adjacent 5’-TC step, which modulates the binding preferences of Asn74 and Gln78. The Gln78 residue interacts with the 3’-AA step of YRE2 on the Crick DNA strand (TTACGT**AA**), which consequently directs Asn74 towards the flanking 5’-TC step to form specific contacts with the cytosine (Figure 2B). The OYE2-environment appears to be less favourable, and contributes to a reduction in strength of Phe81-T interactions compared to YRE2-ATR1 (Figure 2B, Figure S3).

In the case of monomer 2, we observe identical strengths for specific contacts formed by Asn74, Gln78 and Phe81 for the two genomic environments (Figure 2B). Moreover, YRE2 is a palindrome with a central CG step, which allows specific Arg82-G contacts by both monomers (TTA**CG**TAA). Our simulations indicate, however, that only one monomer exhibits strong specific Arg82-G interactions, while the other monomer forms mainly salt bridge contacts between Arg82 and the DNA backbone (Figure 2B), though which monomer depends on the flanking sequence. The similarity of the specific contacts network is surprising in view of the differences in the 3’-flanks, e.g. YRE2 in the ATR1-environment is flanked by the 3’-GC step and in the OYE2-environment – by the 3’-TG step. The ATR1-environment allows for an additional specific Arg70-G contact on the Crick-DNA strand and thus a tighter a positioning of monomer 2 inside the major groove, while the OYE2-environment results in no additional specific contacts. We can trace the observed surprising similarities to the sequence-specific interplay in DNA helical parameters, which we discuss further in section “Helical parameters”.

For YRE3 (TGACAAA), we observe that the two genomic environments, ATR1 and OYE2, allow Yap1 monomer 1 to exhibit similar specific contacts formed by Ala77, Gln78 and Phe81 (Figure 2C, Figure S4). However, in the OYE2-environment, YRE3 is flanked by the 5’-GA step, leading to the specific Arg70-G contact, which allows stronger Asn74-YRE3 and Arg82-YRE3 interactions. In contrast, the adjacent 5’-TA flanking step, in the ATR1-environment, contributes to destabilisation of DNA-monomer 1 interactions. We observe a large standard deviation of the Asn74-YRE3 contact strength, where Asn74 for 20% of the simulation time (Figure S4) completely loses contacts with DNA (Figure 2C). YRE3, which contains TG instead of the TT step (**TG**ACAAA), shows a clear reduction in the specific contacts strength exhibited by Phe81 of monomer 1. However, monomer 1 adjusts to compensate by increasing the strength of the Arg82-G contact compared to YRE1 (Figure 2A, 2C and Figure S2, S4).

We observe that Yap1 monomer 2 forms similar Gln78-YRE3 and Phe81-YRE3 contacts in both genomic environments (Figure 2C and Figure S4). However, in OYE2-environment, YRE3 is flanked by the 3’-AT step (TGACAAA-**AT**), which results in a complete loss of the Asn74-DNA interactions for the Crick strand. The flanking 3’-AT step can potentially create steric clashes between Ala77 and the methyl group of the flanking 5’-T nucleotide on the Crick strand (A**T**-TTTGTCA). This arrangement prevents the deeper placement of monomer 2 inside the major groove, resulting in weaker YRE3-monomer 2 interactions. In contrast, in the ATR1-environment, YRE3 is flanked by the 3’-GT step (or 5’-CA on the Crick strand), which similarly to the 5’-GC step results in specific contacts with Arg70 and Asn74 from the Crick strand (Figure 2C and Figure S4). However, interactions with the flanking 5’-AC step are weaker compared to the 5’-GC step.

Our simulations show that the 5’-GA flanking step is favourable for Yap1-YRE complexation. This, however, is not seen in experimentally derived sequence logos for the majority of BZIP factors, except for Yap8 (which YRE contains extended 5-TGA sites)^9^ and the Maf family (extended 5’-TGC sites),^6^ probably, since sequence logos are generated by looking at each nucleotide position individually, excluding sequence specific effects at the b.p. step level. For Yap1, we observe that Arg70 forms specific contacts with the 5’-GA flanking step, which stabilises the position of Yap1 in the major groove as well as the specific contacts formed between YRE and the five-residues-motif (**N**xx**AQ**xx**FR**). However, our simulations also detect that other adjacent flanking steps provide similar strength of specific Yap1-YRE contacts exploited by the five-residues-motif, including 5’-GC, 5’-CA and 5’-AC steps, suggesting why the flanking sites for Yap1 are not captured by sequence logos. In contrast, comparing Yap1 to Yap8, another member of yeast AP-1 family, we observe modifications within the five-residues-motif (**N**xx**A**QxxFR → **L**xx**S**QxxFR) that can explain the increased preference for extended 5’-TGC sites (**TGA**-TTANNTAA-**TCA**). For instance, highly conserved Asn74 (Yap1) (Rxxx**N**xxAQxxFR) is exchanged for leucine (Leu26, Yap8). The bulkier and more hydrophobic Leu26 helps to reposition Arg22 (Arg70 in Yap1, **R**xxxNxxAQxxFR) to strengthen Arg22-G (of T**G**A) contacts to enhance the recognition of the extended 5’-GA sites. Furthermore, the modification of Ala77 (Yap1, RxxxNxx**A**QxxFR) to a Ser residue (Ser29, Yap8) helps to discriminate between a 5’-GA and 5’-GC step as Ser29 forms specific contacts with the 5’-A nucleotide (TG**A**-TTANNTAA-TCA).^9^

Our simulations show that in comparison to a 5’-GA step, an adjacent 5’-GC step directs the Asn74 residue towards the YRE flanks, which is in particular exploited by the Maf family. Furthermore, the Maf family contains the Ala → Tyr modification within the DNA-recognition motif (Yap1: RxxxNxx**A**QxxFR → Maf: RxxxNxx**Y**AxxCR), which increases the preference for extended 5-TGC sites (**TGC**-TGACGTCA-**GCA**). The Tyr residue stabilises the orientation of the Arg and Asn residues (**R**xxx**N**xxYAxxCR) towards specific contacts with the 5’-GC step (T**GC**-TGACGTCA-GCA). In addition, the Tyr residue forms hydrophobic interactions with the 5’-C nucleotide (TG**C**-TGACGTCA-GCA), which helps to discriminate between a 5’-GC and 5’-GA step.^6^

Our data allows differentiating between high and low affinity response elements flanking sequences. In accordance with a recent HT-SELLEX and bioinformatics study^19^ focusing on the BZIP factors Fos-Jun and NFIL3, we show that the flanking 5’-GA step promotes stable selective DNA-Yap1 binding. The flanking 5’-TA step, which is listed as high affinity for the NFIL3 factor, is less favourable for Yap1; thus, indicating that *at least* two adjacent flanking nucleotides that surrounds the response element can modulate the selectivity among different BZIP factors. We recognise the flanking sites containing a flanking 5’-T nucleotide as low affinity for Yap1, similarly to what was observed for the Fos-Jun and NFIL3 factors. At the same time, we hypothesize that binding of BZIP factors to response elements flanked by 5’-T nucleotides can be beneficial for a cooperative association of multiple transcription factors; the steric hindrance caused by the thymines in the major groove can lower the energy cost for a bending deformation of DNA towards the minor groove facilitating the binding of another transcription factor.^47^

Our simulations show that Yap1 selectivity towards the flanking 5’-GA step can be explained by the specific Arg70-G contact, this direct readout mechanism is insufficient to explain how BZIP factors Jun and NFIL3 that contain an Arg → Lys modification (Jun: **K**xxxNxxAAxxCR, NFIL3: **K**xxxNxxAAxxSR) also show a strong preference for the 5’-GA steps. The shorter side chain of lysine compared to arginine will have more difficulties to approach the major groove for specific contacts with the 5-GA step, suggesting that even for transcription factors where recognition is dominated by the direct readout, the dynamics of DNA could contribute to fine-tuning of the protein-DNA complexation.

### Helical Parameters

DNA sequence-specific dynamics can modulate the recognition by adjusting the helical and groove parameters to resemble the bioactive conformation,^16^ facilitating transcription factors binding. For major groove binders like BZIP factors, shift that involves the relative displacement of b.p. steps in and out of the major groove, and slide that involves the relative displacement of b.p. between the backbones of two DNA strands, are potentially the key parameters that can facilitate the formation of specific protein side chains-DNA bases contacts. Our data show that Yap1 forms a similar number of specific contacts with the three YREs, but the flanking environment impacts the strengths of the protein-DNA contacts. To further understand the molecular mechanism of DNA recognition by Yap1, we analyse the helical parameters for both bound and unbound DNA. Indeed, we observe changes in shift, slide and twist in the presence of Yap1 (see the corresponding distributions, presented in the 5’-> 3’ direction, in Figures 3A-C, S5A-C, S6A-C). Our data also show that the nearest four (shift, slide) to six (twist) flanking nucleotides modulate the helical parameters distributions within the response element for Yap1-bound DNA, which fine-tunes the direct read-out mechanism. In particular, broad shift and slide distributions within the response elements result in the loss or rearrangements of specific contacts.

**Figure 3.**
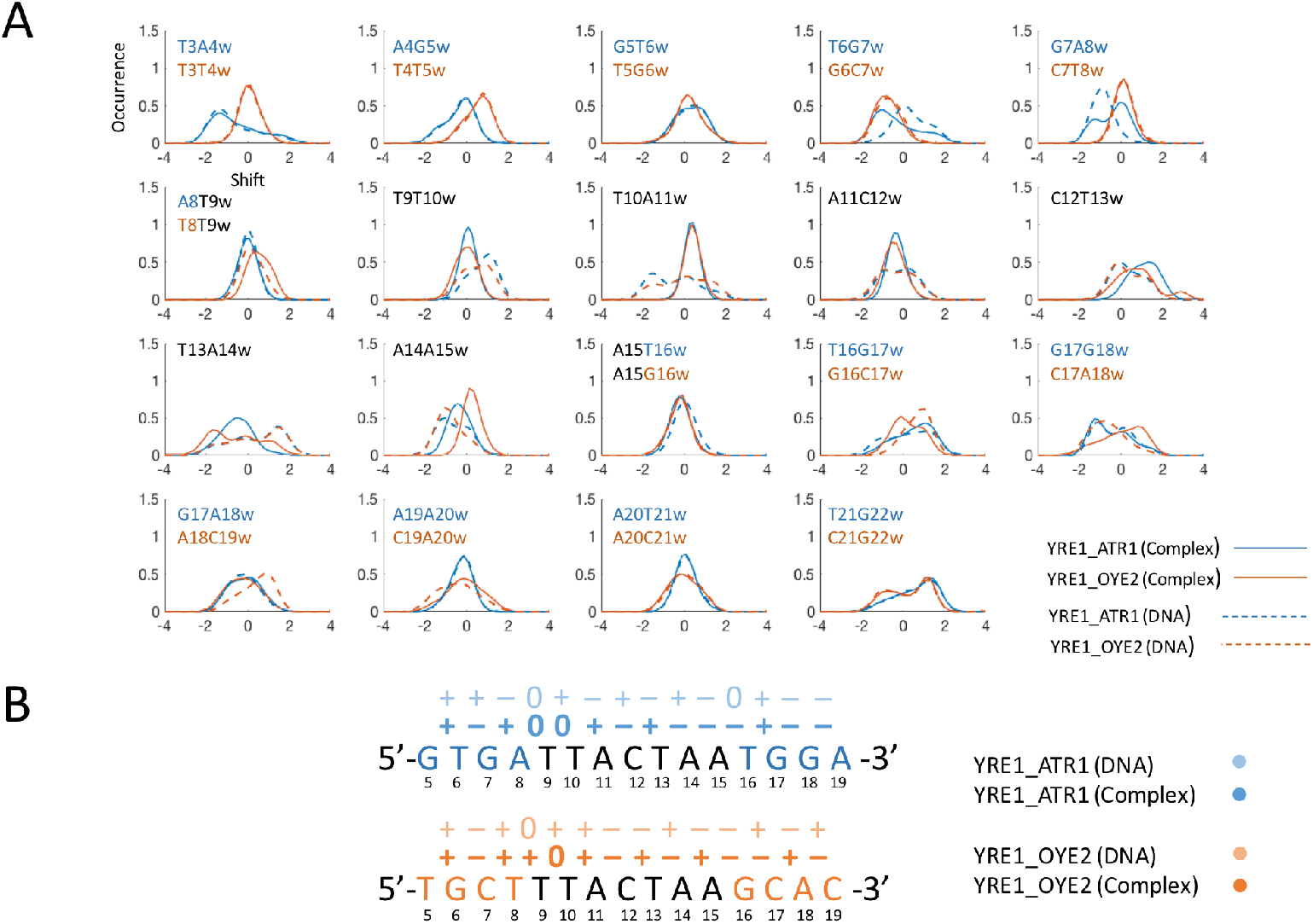
**A.** Normalised shift distributions for unbound (dashed lines) and Yap1-bound (thick lines) YRE1-DNA **B.** Most populated shift (the signs “–”, “+” and “0” indicate negative, positive and neutral shift) for unbound (light colour) and Yap1-bound (dark colour) YRE1-DNA with the four adjacent 5’- and 3’-flanking nucleotides. The genomic environments, ATR1 and OYE2, are coloured blue and orange, respectively.

Analysing shift distributions for YRE1 (direction 5’-> 3’, Figure 3A), we see that in the ATR1-environment, the rearrangement of shift for the 5’-flanking b.p. steps, TG and GA (**TGA**-TTACTAA) in the presence of Yap1, provides an optimal environment for the specific Arg70-G contacts. However, the broad distribution of shift values for the GA step results in a high standard deviation for Arg70-G contact strength, suggesting a highly dynamic nature of this contact (Figure 2A). In the OYE2-environment, the 5’-flanking b.p. steps, GC and CT (**GCT**-TTACTAA) are more rigid and exhibit similar shift distributions for bound and unbound DNA. The stiffer flanking environment together with the positive shift of the TT step (GC**T**-**T**TACTAA) prevents deep association of Yap1 monomer 1 within DNA major groove, resulting in weaker Asn74-YRE interactions.

For Yap1-bound YRE1-DNA, both genomic environments result in similar narrow shift distributions for the first half-site of YRE1 (**TTAC**TAA), which explains the similarly exploited specific contacts by Gln78 and Phe81 of Yap1 monomer 1. The observed negative shift of the AC step allows a more positive shift of the CT step (TTA**CT**AA) step, which favours the Arg82-G interactions (TTA**G**TAA). Though, the CT step shows a broad shift distribution, the positive shift population represents the presence of the Arg82-G contact (~50% for ATR1 and ~30% for OYE2, Figure S2A and S7).

For the second half-site of Yap1-bound YRE1 (TTA**CTAA**), the two genomic environments exhibit more distinct shift distributions, reflecting noticeable differences in YRE1-monomer 2 interactions (Figure 2A). The Y..R (C-**TA**-A) environment makes the TA step of the second half-site more dynamic, compared to the first half-site in the Y..Y (T-**TA**-C) environment. This contributes to broad shift distributions of the TA and AA steps, which triggers the conformational change of Yap1 monomer 2, where the α-helix slides along the major groove to preserve strong Phe81 contacts with the TT step on the Crick strand (**TT**AGTAA). This in turn repositions Gln78 towards the outer AA step (TTACT**AA**), which consequently reduces the specific contacts of Asn74 with the AA step (Figure S2A). This conformational change is less pronounced for the OYE2-environment, where YRE1 is stabilised by the GC and CA 3’-flanking steps (TTACTAA-**GCA**), which alter they shift to favour the Asn74-C and Arg70-G interactions on the Crick strand (T**GC**-TTAGTAA). The combination of these interactions and a slight positive average shift of the AA step stabilise the orientation of Yap1 monomer 2 towards DNA major groove.

For palindromic YRE2, the ATR1-environment constitutes similar 5’-flanking sites as YRE1_ATR1. Analysing shift distributions (Figure 4A), we observe similar shift distributions and the specific contacts network within the first YRE2 half-site (**TTAC**GTAA), involving residues Asn74, Gln78 and Phe81. In contrast, in the OYE2-environment, the 5’-flanking sites (**ATC**-TTACGTAA) limit Yap1 to restrain shift values within YRE2, resulting in broad distributions for the TA and AC steps (T**TAC**GTAA), leading to Yap1 monomer 1 exhibiting somewhat different contacts with YRE2 in the OYE2-environment compared to the ATR1-environment (Figure 2B). The TC and CT steps (A**TC**-**T**TACGTAA) adjust shift to favour the Asn74-C specific contacts, which in turn rearranges shift of the TT step (**TT**ACGTAA) leading to broad shift distributions of the TA and AC steps (T**TAC**GTAA). The shift frustrations explain the weaker Phe81-T contacts and the repositioning of Gln78 towards the AA step on the Crick strand (Figure 2B).

**Figure 4.**
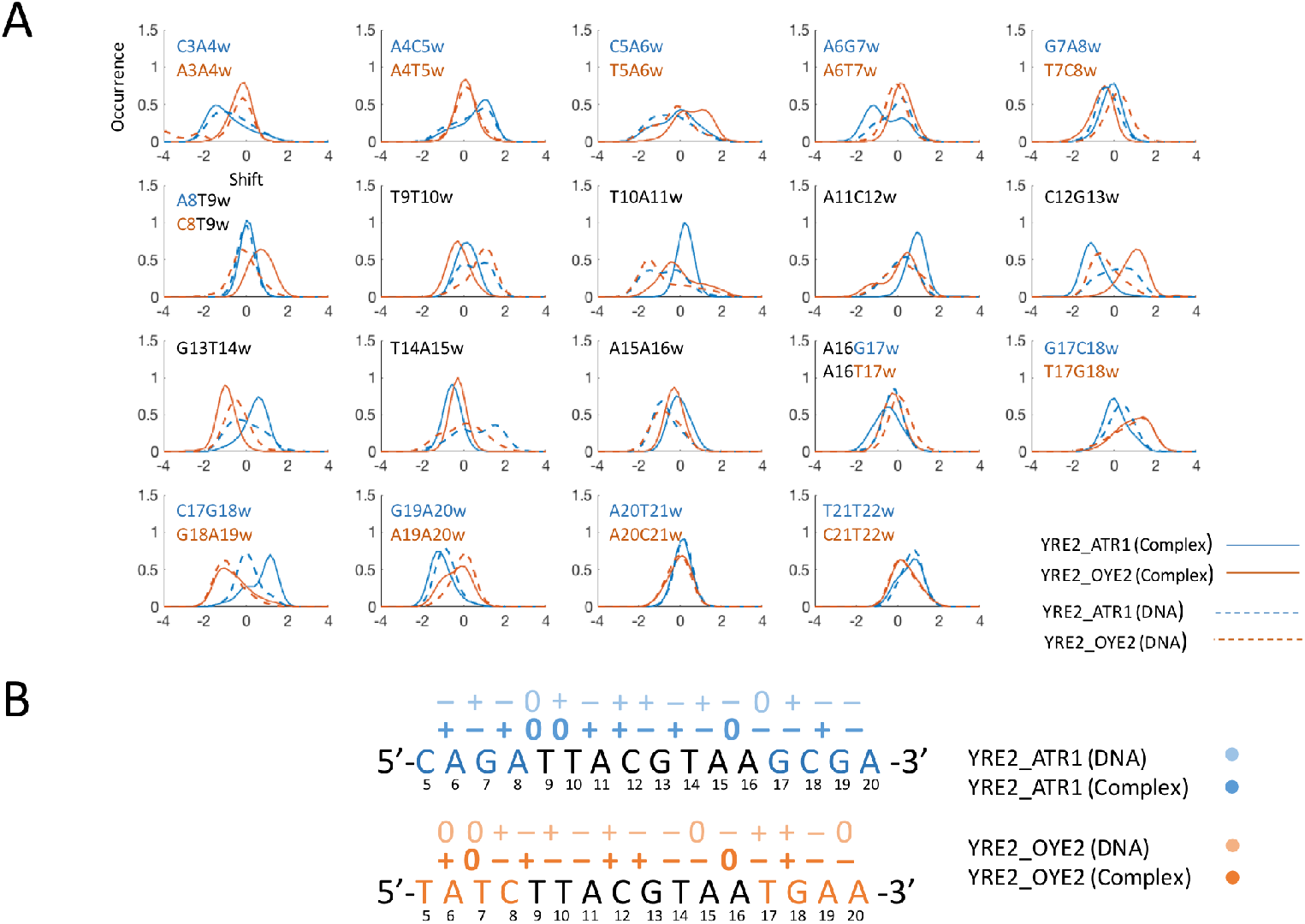
**A.** Normalised shift distributions for unbound (dashed lines) and Yap1-bound (thick lines) YRE2-DNA **B.** Most populated shift (the signs “–”, “+” and “0” indicate negative, positive and neutral shift) for unbound (light colour) and Yap1-bound (dark colour) YRE2-DNA with the four adjacent 5’- and 3’-flanking nucleotides. The genomic environments, ATR1 and OYE2, are coloured blue and orange respectively.

The shift of the CG and GT steps (TTA**CGT**AA) determine which Yap1 monomer will exhibit the specific Arg82-G contacts. In the ATR1-environment, the shift values of the CG and GT steps in Yap1-bound DNA create a suitable environment for Arg82(2)-G_“w”_ interactions and weaker Arg82(1)-G_“c”_ interactions (indices “w” and “c” indicate Watson and Crick DNA strands, respectively). The reverse trend is observed for the OYE2-environment. The 3’-flanking sites of both genomic environments contribute to similar shift distributions for the TA and AA steps (TTACG**TAA**), resulting in the identical contact network for Yap1 monomer 2 with the TAA trinucleotide (Figure 2B).

For YRE3, the OYE2-environment contains a similar 5’-flanking sequence as YRE1_ATR1 and YRE2_ATR1, with a GA step coupled to a soft CG step (**CGA**-TGACAAA). Similarly, for Yap1-bound DNA (Figure 5A), the shift values of the CG and GA steps (**CGA**-TGACAAA) stabilize the specific Arg70-G contact. This flanking environment also allows Yap1 monomer 1 to adjust the shift distributions within the first half-site of YRE3 (**TGAC**AAA), which contributes to stable specific contacts with Asn74, Gln78, Phe81 and Arg82. Furthermore, by reducing the shift of the TG step (**TG**ACAAA) that allows increasing the strength of the Phe81-T nonspecific contacts (Figure S4B), YRE3 compensates the missing TT dinucleotide at the beginning of YRE and the weaker specific Phe81-T interactions. In the ATR1-environment, the 5’-flanking site is more rigid (**ATA**-TGACAAA). This restricts shift adjustments for the TG and GA steps (**TGA**CAAA) in the presence of Yap1. The broad shift distributions induce sliding and bending of Yap1 monomer 1 basic region to maintain the Phe81-T interactions, which in turn increases fluctuations of the Asn74-YRE3 contacts (Figure 2C and S4A). In both genomic environments, in Yap1-bound DNA, the shift values of the central AC and CA steps provide a suitable environment for the specific Arg82-G interactions from the Crick strand side.

**Figure 5.**
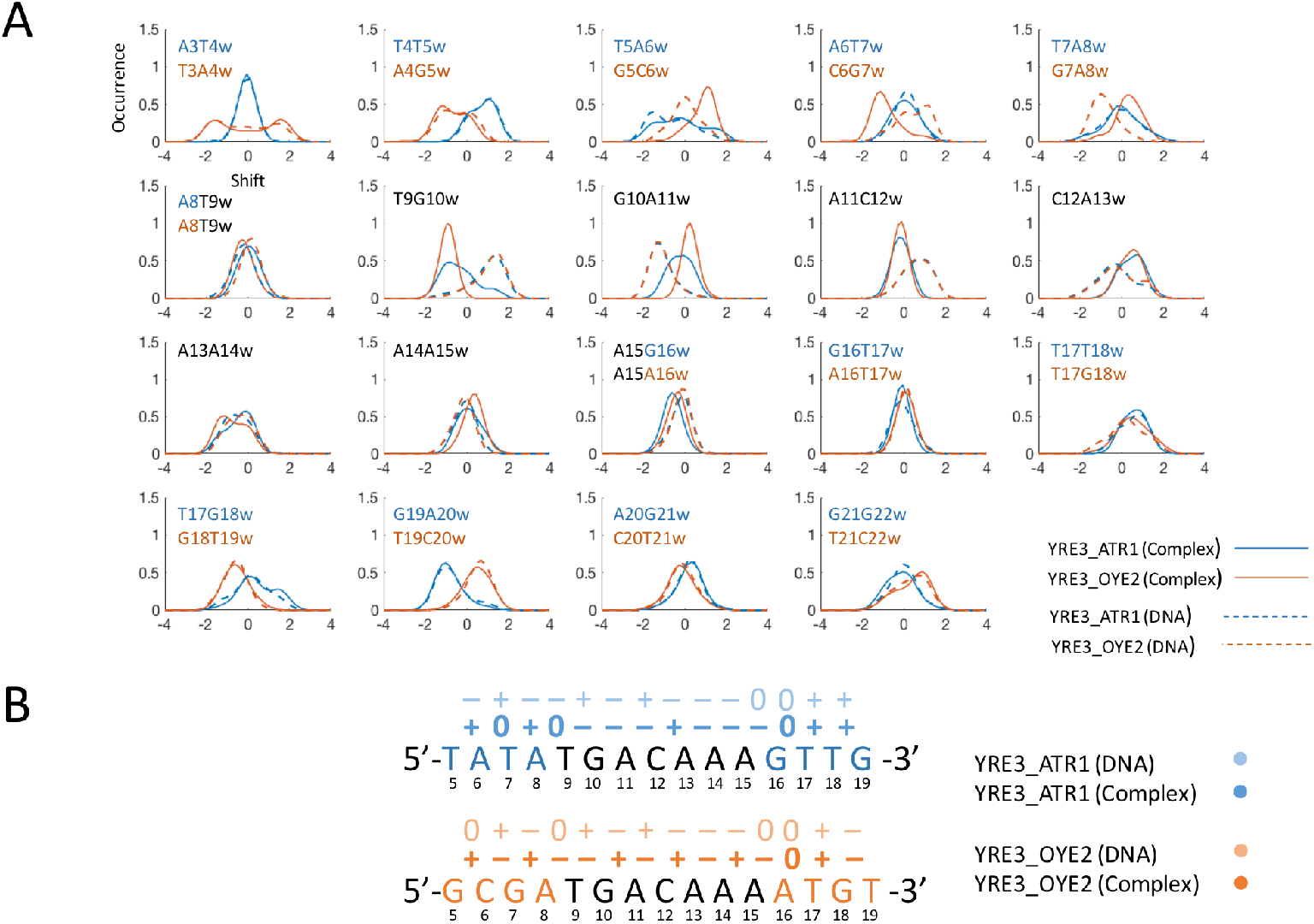
**A.** Normalised shift distributions for unbound (dashed lines) and Yap1-bound (thick lines) YRE3-DNA **B.** Most populated shift (the signs “–”, “+” and “0” indicate negative, positive and neutral shift) for unbound (light colour) and Yap1-bound (dark colour) YRE3-DNA with the four adjacent 5’- and 3’-flanking nucleotides. The genomic environments, ATR1 and OYE2, are coloured blue and orange respectively.

In both genomic environments, for the second half-site of YRE3 (TGA**CAAA**) that interacts with Yap1 monomer 2, we observe no major alterations of shift distributions for Yap1-bound DNA with respect to unbound DNA (Figure 5A). However, the thymines on the Crick strand (**TTT**GTCA) result in either large fluctuations (ATR1) or loss (OYE2) of specific contacts exploited by Asn74 (Figure 2C). In the OYE2-environment, the presence of a stiff 3’-AT flanking step contributes to a negative shift of the extended AA step (TGACAA**A-A**T), allowing for an additional hydrophobic contact, Ala77-T (A**T**-TTTGTCA) on the Crick strand. This contact acts as a steric barrier, preventing the deeper placement of Yap1 monomer 2 within the major groove, thus explaining the loss of Asn74-YRE3 interactions. In the ATR1-environment, YRE3 is stabilized by an adjacent 3’-GT flanking step (TGACAAA-**GT**), where the shift values of the flanking AG and GT steps (TGACAA**A-GT**) provide Asn74-C and Arg70-A interactions with the AC step on the Crick strand (A**C**-TTTGTCA) for ~40% of the trajectory (Figure 2C and S4A).

Moreover, changes in shift are coupled to changes in twist through DNA backbone BI → BII transitions;^14,48,49^ therefore, we also observe an alteration of twist for Yap1-bound DNA. This observation is interesting in the context of DNA supercoiling and its role in eukaryotic transcriptional control.^50^ Together with writhe, twist modulates DNA supercoiling transitions along the chromatin fibre. Recently, we have shown that MafB (a member of human BZIP family)^11^ asymmetrically changes the sequence-specific response of DNA to torsional stress, making DNA effectively more rigid. The molecular mechanism of the observed phenomenon is based upon MafB forming a number of specific contacts with the torsionally flexible YR dinucleotide steps thus restraining their shift. Instead more torsionally rigid dinucleotides are forced to absorb the imposed torsion, resulting in an increased energy cost of twisting in comparison with unbound DNA. Therefore, the degree to which shift, and to less extend slide, can adjust upon association with transcription factors will locally modulate the torsional rigidity of DNA. We hypothesise that the increased local torsional rigidity of DNA impacts how long promoters stay open for transcription, hence regulating transcription initiation rates.

### Ion populations

The flanking sites differences are also reflected in K+ ion populations within the major and minor DNA grooves, which persist in the protein-bound state (Figure S8-S11). This is interesting from a perspective of random N-terminal tails interactions with DNA, in particular in the flanking regions. The majority of BZIP factors, including Yap1, have substantial random coil N-terminal tails, rich in positively charged residues.^9^ Thus, the observed differences in ion populations could potentially fine-tune the DNA-residence time of the protein. In addition, from our simulations, we observe that an adjacent 5’-GA step provides the more stable Arg70-G interactions compared to a 5’-GC step, even though both flanking steps undergo similar alterations in shift for Yap1-bound DNA to set the environment for the Arg70-contact. Analysing ion-distributions (Figure S8), we observe for unbound DNA, consistent with the previous findings,^51^ that a GC step accumulates a higher K+ population in the major groove compared to a GA step. The K+ population around the GC step is not completely depleted in the presence of Yap1, resulting in a competition with the Arg70-G specific contact. We observe similar response for a GT step, thus explaining the large fluctuations of the Arg70-A interactions with the AC step in YRE3_OYE2.

### Bioinformatics analysis of model genes expression levels

To verify our hypothesis, we analyse the differential expression level of ATR1 and OYE2 genes of two yeast strains, wild type versus *yap1Δ*, over time under sodium selenite stress that activates Yap1 (Salin *et al*., BMC Genomics 2008).^44^ We see that the ATR1 gene is significantly more differentially expressed compared to OYE2 (Figure 6). In the ATR1-environment, the Yap1 response elements, YRE1 and YRE2, are surrounded by the contacts-favouring 5’-GA flanking sites (RYGA-YRE1 and YRGA-YRE2), whereas in the OYE2-environment, only one of the response elements, YRE3, contains this 5’-flanking motif (RYGA-YRE3). In addition, in the OYE2-environment, two of the response elements, YRE1 and YRE3, are surrounded by an adjacent 5’-T nucleotide (for YRE1, Watson strand: **T**TTACTAA; for YRE3, Crick strand: **T**TTTGTCA), which due to a steric hindrance, result in a reduced number of specific contacts. Nevertheless, our simulations imply that adjacent 5’-T flanking nucleotides will not completely abolish Yap1 binding; instead we believe this flanking site can be an important motif for a cooperative binding of transcription factors, if the complementary DNA strand contains a more favourable flanking environment. For instance, for YRE3 the OYE2-environment contains a favourable 5’-flanking environment on the Watson strand (5’-**GCGA-**TGACAAA), which stabilizes contacts between Yap1 monomer 1 and the first YRE3 half-site (**TGAC**AAA). The other YRE3 half-site (**TTTG**TCA, Crick strand) that interacts with Yap1 monomer 2 contains an extended region of thymines (5’-AC**AT-TTT**GTCA), which sterically restricts the deeper association of monomer 2 with DNA major groove, thus reducing the strength of specific protein-DNA contacts. This can, depending on the 3D-architecture of the Yeast genome, induce a more favourable topologic environment to enhance Yap1 binding to the other two response elements, YRE1 and YRE2, which are located about 50 nucleotides from YRE3_OYE2 (Figure S1).

**Figure 6.**
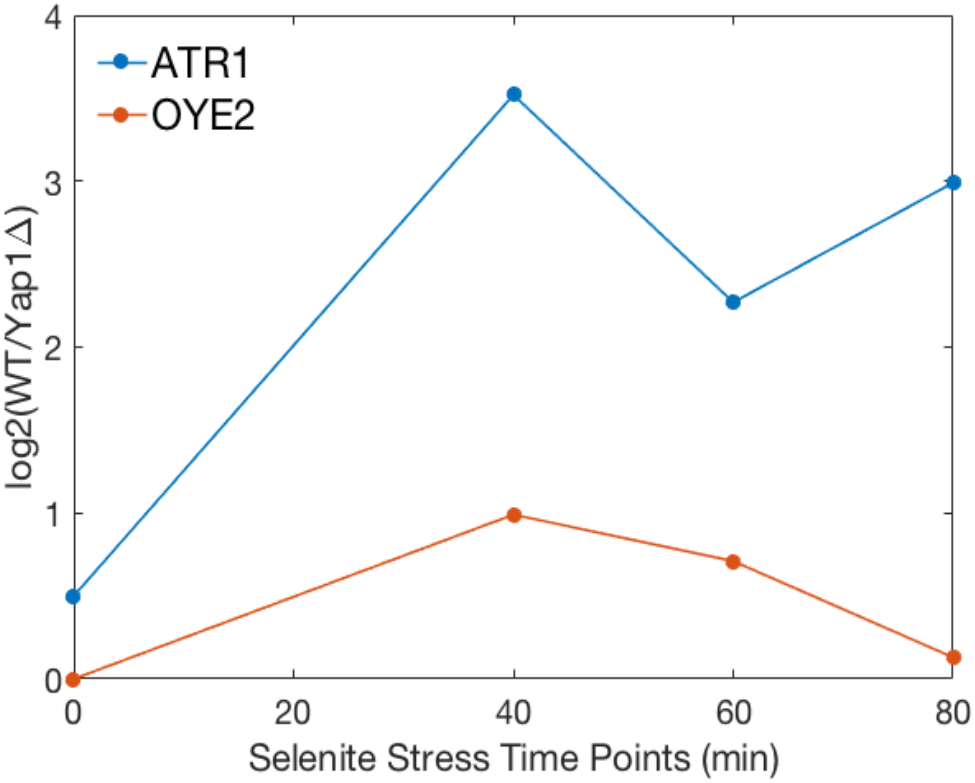
Differential expression levels of Yap1 target genes, ATR1 and OYE2, under selenite stress. Differential expression level between wild type cells (WT) and cells missing yap1 gene (Yap1Δ) were obtained from MicroArray quantification at different time points after selenite stress induction.

To further investigate the role of YREs flanking environment in the regulation of gene expression, we use a library of 100 million randomly generated promoters of 80 b.p. length and their associated expression levels, given by a dual reporter gene system.^45^ The expression levels have been divided into 18 expression bins from the weakest expression (0) to the strongest (17). We filter out the sequences containing only one YRE per promoter sequence, with the four adjacent 5’-flanking and 3’-flanking nucleotides corresponding to the ATR1- and the OYE2-environments, and check the relative expression level. Our analysis shows that YRE1 provides a stronger expression response in the ATR1-environment than in the OYE2-environment, 7 versus 5 in arbitrary expression units. This is consistent with the discussed preference for the flanking 5’-RYGA sequence present in the ATR1-environment. YRE2 provides similar associated expression levels around 8 in arbitrary expression units, however, irrespective of the flanking environment. YRE3 was only found with the flanking sequences resembling, but not identical to the ATR1-environment. The associated expression levels of YRE3-like model promoters vary significantly, with an average expression level of 9 and a standard deviation of 5.7. The extracted expression levels for YRE2 and YRE3 do not show a clear correlation to our predictions about the flanking sites and the strength of binding. One possible explanation is that Yap1 contains an 80-residues long N-terminal tail rich in positively charged residues, which was not included in the model but which could stabilise the protein-DNA interactions. Also, we cannot exclude the possibility that the extracted expression levels also depend on other factors, such as cooperative action between several transcription factors and competitive binding by other proteins, etc. In addition, our simulations show that the palindromic nature of YRE2 (TTA**CG**TAA) allows Yap1 to form more specific contacts with DNA, which could explain the lesser sensitivity towards the flanking environments. We also observe for YRE2 that the less favourable 5’-flanking sites in the OYE2-environment result in the rearrangement of Yap1-DNA specific contacts rather than a reduction; this can also contribute to similar expression levels. For YRE3, the high standard deviation in expression levels of the resembling ATR1-environments may suggest an impact of longer than four nucleotides flanking sites. However, based on our simulations we believe that YRE3 can be a cooperative driven/dependent Yap1 response element. Yap1 bound to YRE3, as discussed, shows a reduction of Phe81-YRE3 hydrophobic contacts, due to the presence of the TG step (**TG**ACAAA) instead of a more favourable TT step. Thus, compared to binding of Yap1 to YRE1 and YRE2, the weaker Yap1 binding to YRE3 may induce a different dynamics of DNA, such as bending towards the minor groove, which could effectively enhance the cooperative binding of transcription factors that can result in an overall stronger transcription response. Hence, the YRE3_ATR1 resembling promoters that show lower expression levels might exhibit weaker cooperative actions between transcription factors.

## Conclusion

AP-1 BZIP transcription factors execute their gene regulatory programs through specific binding to the corresponding DNA response elements. The DNA recognition mechanism of AP-1 proteins follows the direct read-out principle, when the highly conserved, among the subfamilies, motifs of the protein basic region forms specific contacts with the bases of the response elements. The response elements recognised by AP-1 vary both in lengths, seven-14 b.p., and the nucleotide composition.^3,5,6^ Moreover, these short specific DNA stretches occur in organisms’ genomes much more frequently than the actual genes regulated by the transcription factors, which implies that the complexity of the recognition process goes beyond the ‘simplistic’ direct read-out mechanism. Using microsecond long all-atom molecular dynamics simulations, we addressed two additional aspects: the roles of DNA sequence-specific dynamics and the response elements’ flanking sequences, for the molecular recognition process, protein-DNA complex stability, and in general for the regulation of DNA transcription reaction. As a model system, we employed yeast Yap1 basic leucine zipper transcription factor interacting with three different Yap1 response elements (YRE1: TTACTAA, YRE2: TTACGTAA, YRE3: TGACAAA) from two native genomic environments (ATR1 and OYE2).

Our results show that for the recognition by Yap1, DNA sequence-specific dynamics fine-tunes the direct readout mechanism. Adjustment of DNA base pairs shift (and indirectly twist) and slide helical parameters within the response element creates an optimal environment in the major groove to allow for stable, specific contacts with the five-residues-motif of Yap1 (**N**xxx**AQ**xx**FR**). Combining our observations with the analysis of the available crystallographic structures of BZIP-DNA complexes (Figure S12) we conclude: BZIP factors favour a specific shift environment within the response element that enables specific protein-DNA contacts. Previous MD studies^16^ confirm that DNA dynamics facilitates the conformational transition of unbound DNA to its bioactive state; where the transition proceeds more smoothly if the DNA sequence is the corresponding consensus sequence for a particular protein, rather than a random sequence. Our simulations further show that this conformational transition also depends on the flanking sequence environment, surrounding the response element. We observe that four to six flanking nucleotides impact how efficiently Yap1 can restrain shift distributions of the b.p. steps within the response element. Unfavourable flanking sites result in broad shift distributions within YREs, which either causes a reduction or a rearrangement of specific contacts exploited by Yap1. This, we suggest, will influence the binding affinity and allow the transcription factor to discriminate between different genomic locations containing the same response element. In addition, we know from the previous studies^14,48,49^ that change in shift brings changes in twist, important for the regulation of DNA supercoiling transitions and, by extension, transcriptional control. Thus, the level of adjustment of b.p. shift when a transcription factor is bound to DNA, directly translates into DNA local twist flexibility, and consequently the energetic cost of DNA supercoiling transitions.^11^ The described molecular mechanism, we believe, allows the transcription factor to regulate the opening of gene promoters and subsequently their firing potentials, and by extension, the gene expression levels.

## Acknowledgement

This work was supported by Swedish Foundation for Strategic Research SSF Grant [ITM17-0431] and Hasselblad Foundation Prize to A.R. The authors thank Swedish National Infrastructure for Computing (SNIC) for the generous provision of computing resources.

## References

1. Shaulian, E. & Karin, M. AP-1 as a regulator of cell life and death. Nat. Cell Biol. 4, E131–E136 (2002).

2. Garces de Los Fayos Alonso, I. et al. The Role of Activator Protein-1 (AP-1) Family Members in CD30-Positive Lymphomas. Cancers (Basel). 10, 93 (2018).

3. Fujii, Y., Shimizu, T., Toda, T., Yanagida, M. & Hakoshima, T. Structural basis for the diversity of DNA recognition by bZIP transcription factors. Nat. Struct. Biol. 7, 889 (2000).

4. Karin, M., Liu, Z. & Zandi, E. AP-1 function and regulation. Curr. Opin. Cell Biol. 9, 240–246 (1997).

5. Rodríguez-Martínez, J. A., Reinke, A. W., Bhimsaria, D., Keating, A. E. & Ansari, A. Z. Combinatorial bZIP dimers display complex DNA-binding specificity landscapes. Elife 6, e19272 (2017).

6. Kurokawa, H. et al. Structural basis of alternative DNA recognition by Maf transcription factors. Mol. Cell. Biol. 29, 6232–6244 (2009).

7. Rodrigues-Pousada, C. et al. Yeast AP-1 like transcription factors (Yap) and stress response: a current overview. Microb. cell (Graz, Austria) 6, 267–285 (2019).

8. Amaral, C. et al. Two Residues in the Basic Region of the Yeast Transcription Factor Yap8 Are Crucial for Its DNA-Binding Specificity. PLoS One 8, e83328 (2013).

9. Maciaszczyk-Dziubinska, E. et al. The ancillary N-terminal region of the yeast AP-1 transcription factor Yap8 contributes to its DNA binding specificity. Nucleic Acids Res. (2020). doi:10.1093/nar/gkaa316

10. Sitlani, A. & Crothers, D. M. Fos and Jun do not bend the AP-1 recognition site. Proc. Natl. Acad. Sci. U. S. A. 93, 3248–3252 (1996).

11. Hörberg, J. & Reymer, A. BZIP Transcription Factors Modulate DNA Supercoiling Transitions. bioRxiv 2019.12.13.875146 (2019). doi:10.1101/2019.12.13.875146

12. Olson, W. K., Gorin, A. A., Lu, X.-J. J., Hock, L. M. & Zhurkin, V. B. DNA sequence-dependent deformability deduced from protein-DNA crystal complexes. Proc. Natl. Acad. Sci. U. S. A. 95, 11163–11168 (1998).

13. Dans, P. D., Pérez, A., Faustino, I., Lavery, R. & Orozco, M. Exploring polymorphisms in B-DNA helical conformations. Nucleic Acids Res. 40, 10668–78 (2012).

14. Pasi, M. et al. μABC: a systematic microsecond molecular dynamics study of tetranucleotide sequence effects in B-DNA. Nucleic Acids Res. 42, 12272–12283 (2014).

15. Balaceanu, A. et al. Modulation of the helical properties of DNA: next-to-nearest neighbour effects and beyond. Nucleic Acids Res. 1–13 (2019). doi:10.1093/nar/gkz255

16. Battistini, F. et al. How B-DNA Dynamics Decipher Sequence-Selective Protein Recognition. J. Mol. Biol. 431, 3845–3859 (2019).

17. Dror, I., Rohs, R. & Mandel-Gutfreund, Y. How motif environment influences transcription factor search dynamics: Finding a needle in a haystack. Bioessays 38, 605–612 (2016).

18. Cohen, D. M., Lim, H. W., Won, K. J. & Steger, D. J. Shared nucleotide flanks confer transcriptional competency to bZip core motifs. Nucleic Acids Res. 46, 8371–8384 (2018).

19. Yella, V. R. et al. Flexibility and structure of flanking DNA impact transcription factor affinity for its core motif. Nucleic Acids Res. 46, 11883–11897 (2018).

20. Nguyên, D.-T., Alarco, A.-M. & Raymond, M. Multiple Yap1p-binding Sites Mediate Induction of the Yeast Major Facilitator FLR1 Gene in Response to Drugs, Oxidants, and Alkylating Agents. J. Biol. Chem. 276, 1138–1145 (2001).

21. Teixeira, M. C. et al. Refining current knowledge on the yeast FLR1 regulatory network by combined experimental and computational approaches. Mol. Biosyst. 6, 2471 (2010).

22. Kuo, D. et al. Coevolution within a transcriptional network by compensatory trans and cis mutations. Genome Res. 20, 1672–1678 (2010).

23. Goudot, C., Etchebest, C., Devaux, F. & Lelandais, G. The Reconstruction of Condition-Specific Transcriptional Modules Provides New Insights in the Evolution of Yeast AP-1 Proteins. PLoS One 6, e20924 (2011).

24. He, X.-J. & Fassler, J. S. Identification of novel Yap1p and Skn7p binding sites involved in the oxidative stress response of Saccharomyces cerevisiae. Mol. Microbiol. 58, 1454–1467 (2005).

25. Tan, K. et al. A systems approach to delineate functions of paralogous transcription factors: role of the Yap family in the DNA damage response. Proc. Natl. Acad. Sci. U. S. A. 105, 2934–2939 (2008).

26. Coordinators, N. R. Database resources of the National Center for Biotechnology Information. Nucleic Acids Res. 46, D8–D13 (2018).

27. Krieger, E. & Vriend, G. YASARA View – molecular graphics for all devices – from smartphones to workstations. Bioinformatics 30, 2981–2982 (2014).

28. The UniProt Consortium. UniProt: the universal protein knowledgebase. Nucleic Acids Res. 45, D158–D169 (2016).

29. Yan, Y., Zhang, D., Zhou, P., Li, B. & Huang, S.-Y. HDOCK: a web server for protein– protein and protein–DNA/RNA docking based on a hybrid strategy. Nucleic Acids Res. 45, W365–W373 (2017).

30. Lavery, R., Zakrzewska, K. & Sklenar, H. JUMNA (junction minimisation of nucleic acids). Comput. Phys. Commun. 91, 135–158 (1995).

31. Abraham, M. J. et al. GROMACS: High performance molecular simulations through multi-level parallelism from laptops to supercomputers. SoftwareX 1–2, 19–25 (2015).

32. Maier, J. A. et al. ff14SB: Improving the Accuracy of Protein Side Chain and Backbone Parameters from ff99SB. J. Chem. Theory Comput. 11, 3696–3713 (2015).

33. Ivani, I. et al. Parmbsc1: a refined force field for DNA simulations. Nat. Methods 13, 55–58 (2016).

34. Mark, P. & Nilsson, L. Structure and Dynamics of the TIP3P, SPC, and SPC/E Water Models at 298 K. J. Phys. Chem. A 105, 9954–9960 (2001).

35. Berendsen, H. J. C., Postma, J. P. M., van Gunsteren, W. F., DiNola, A. & Haak, J. R. Molecular dynamics with coupling to an external bath. J. Chem. Phys. 81, 3684–3690 (1984).

36. Parrinello, M. & Rahman, A. Polymorphic transitions in single crystals: A new molecular dynamics method. J. Appl. Phys. 52, 7182–7190 (1981).

37. Hess, B., Bekker, H., Berendsen, H. J. C. & Fraaije, J. G. E. M. LINCS: A linear constraint solver for molecular simulations. J. Comput. Chem. 18, 1463–1472 (1997).

38. Darden, T., York, D. & Pedersen, L. Particle mesh Ewald: An N⋅log(N) method for Ewald sums in large systems. J. Chem. Phys. 98, 10089–10092 (1993).

39. Páll, S. & Hess, B. A flexible algorithm for calculating pair interactions on SIMD architectures. Comput. Phys. Commun. 184, 2641–2650 (2013).

40. Harvey, S. C., Tan, R. K.-Z. & Cheatham III, T. E. The flying ice cube: Velocity rescaling in molecular dynamics leads to violation of energy equipartition. J. Comput. Chem. 19, 726–740 (1998).

41. Roe, D. R. & Cheatham, T. E. PTRAJ and CPPTRAJ: Software for Processing and Analysis of Molecular Dynamics Trajectory Data. J. Chem. Theory Comput. 9, 3084–3095 (2013).

42. Lavery, R., Moakher, M., Maddocks, J. H., Petkeviciute, D. & Zakrzewska, K. Conformational analysis of nucleic acids revisited: Curves+. Nucleic Acids Res. 37, 5917–5929 (2009).

43. Berman, H. M. et al. The Protein Data Bank. Nucleic Acids Res. 28, 235–242 (2000).

44. Salin, H. et al. Structure and properties of transcriptional networks driving selenite stress response in yeasts. BMC Genomics 9, 333 (2008).

45. de Boer, C. G. et al. Deciphering eukaryotic gene-regulatory logic with 100 million random promoters. Nat. Biotechnol. 38, 56–65 (2020).

46. Pettersen, E. F. et al. UCSF Chimera—A visualization system for exploratory research and analysis. J. Comput. Chem. 25, 1605–1612 (2004).

47. Panne, D., Maniatis, T. & Harrison, S. C. Crystal structure of ATF-2/c-Jun and IRF-3 bound to the interferon-β enhancer. EMBO J. 23, 4384–4393 (2004).

48. Reymer, A., Zakrzewska, K. & Lavery, R. Sequence-dependent response of DNA to torsional stress: a potential biological regulation mechanism. Nucleic Acids Res. 46, 1684–1694 (2018).

49. Hörberg, J. & Reymer, A. A sequence environment modulates the impact of methylation on the torsional rigidity of DNA. Chem. Commun. 54, 11885–11888 (2018).

50. Lavelle, C. DNA torsional stress propagates through chromatin fiber and participates in transcriptional regulation. Nat. Struct. Mol. Biol. 15, 123–5 (2008).

51. Pasi, M., Maddocks, J. H. & Lavery, R. Analyzing ion distributions around DNA: sequence-dependence of potassium ion distributions from microsecond molecular dynamics. Nucleic Acids Res. 43, 2412–2423 (2015).

