## Supplementary figures and tables for "Sequence-specific dynamics of DNA response elements and their flanking sites regulate the recognition by AP-1 transcription factors"


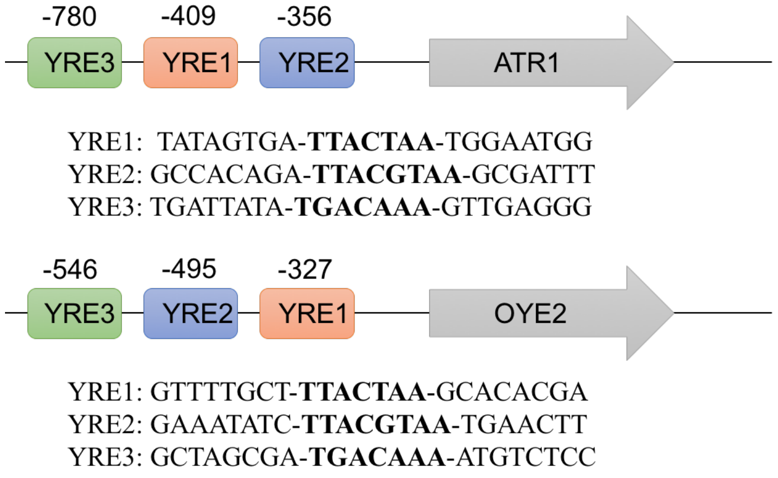


*Figure S1:* The three different Yap1 Response Elements (YREs): YRE1 – TTACTAA, YRE2 – TTACGTAA, and YRE3 – TGACAAA, located in the vicinity of the Yap1 regulated genes, ATR1 and OYE2.

*Table S1:* Parameters used for the homology modelling of Yap1 in YASARA.

| Homology Modelling Parameters | |
|---|---|
| Modelling speed (slow=best) | Slow |
| Number of PSI-BLAST iterations | 6 |
| Maximum number of templates to be used | 1 |
| Maximum number of templates with same sequence | 1 |
| Maximum oligomerization state | 4 |
| Maximum number of alignment variations per template | 5 |
| Maximum number of conformations tried per loop | 50 |
| Maximum number of residues added to the termini | 10 |

A

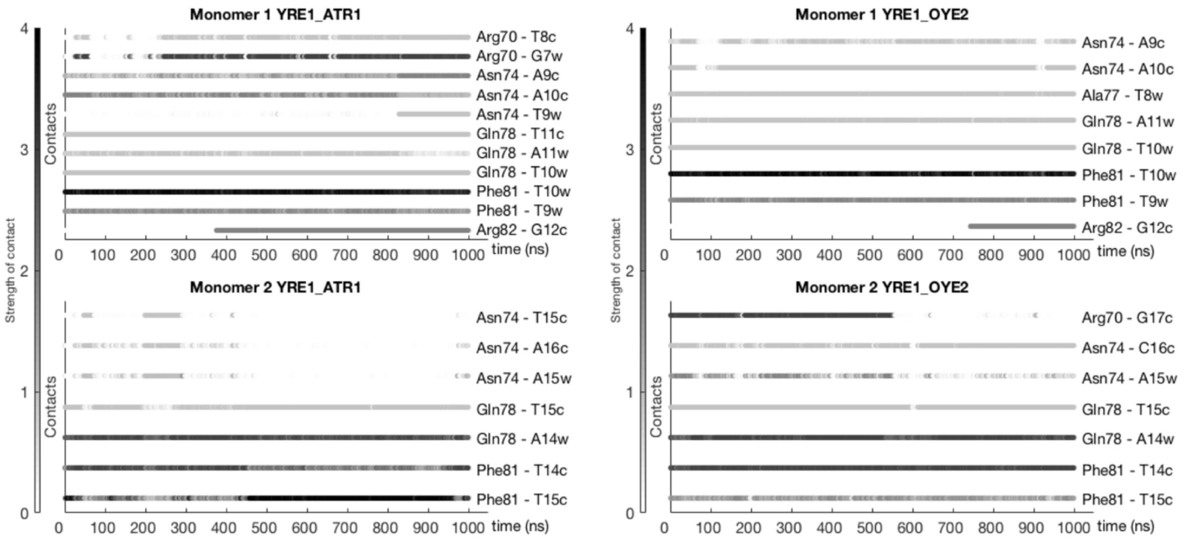


B

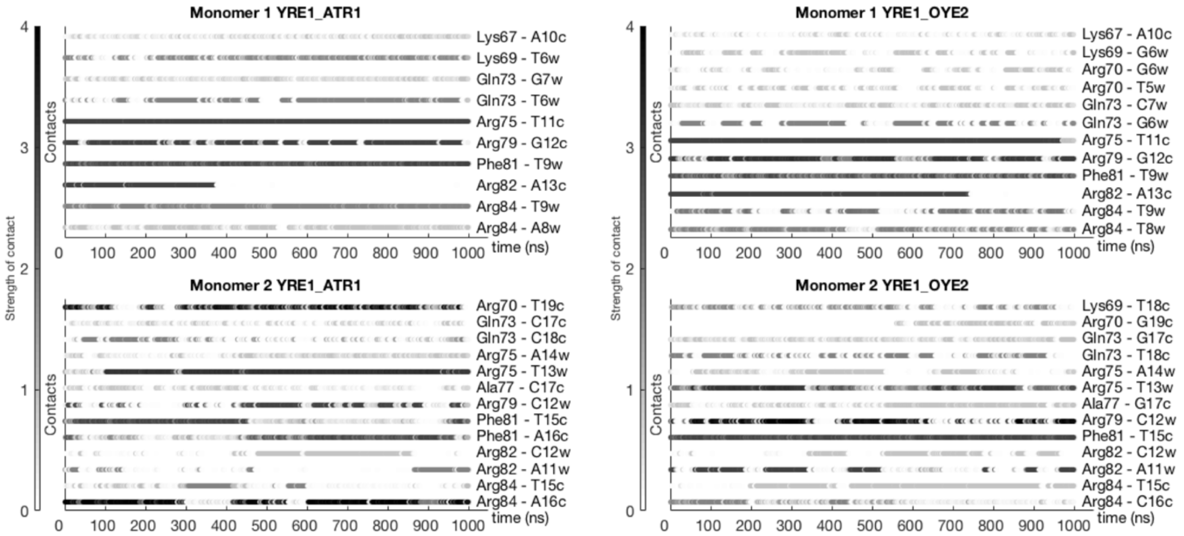


*Figure S2:* Dynamic interaction maps for Yap1 in complex with YRE1 in the ATR1- and OYE2-environments. The maps illustrate the strenght of **A.** specific and **B.** nonspecific Yap1-DNA contacts as a function of time. Indices “w” and “c” reffer to Watson- and Crick-DNA strands.

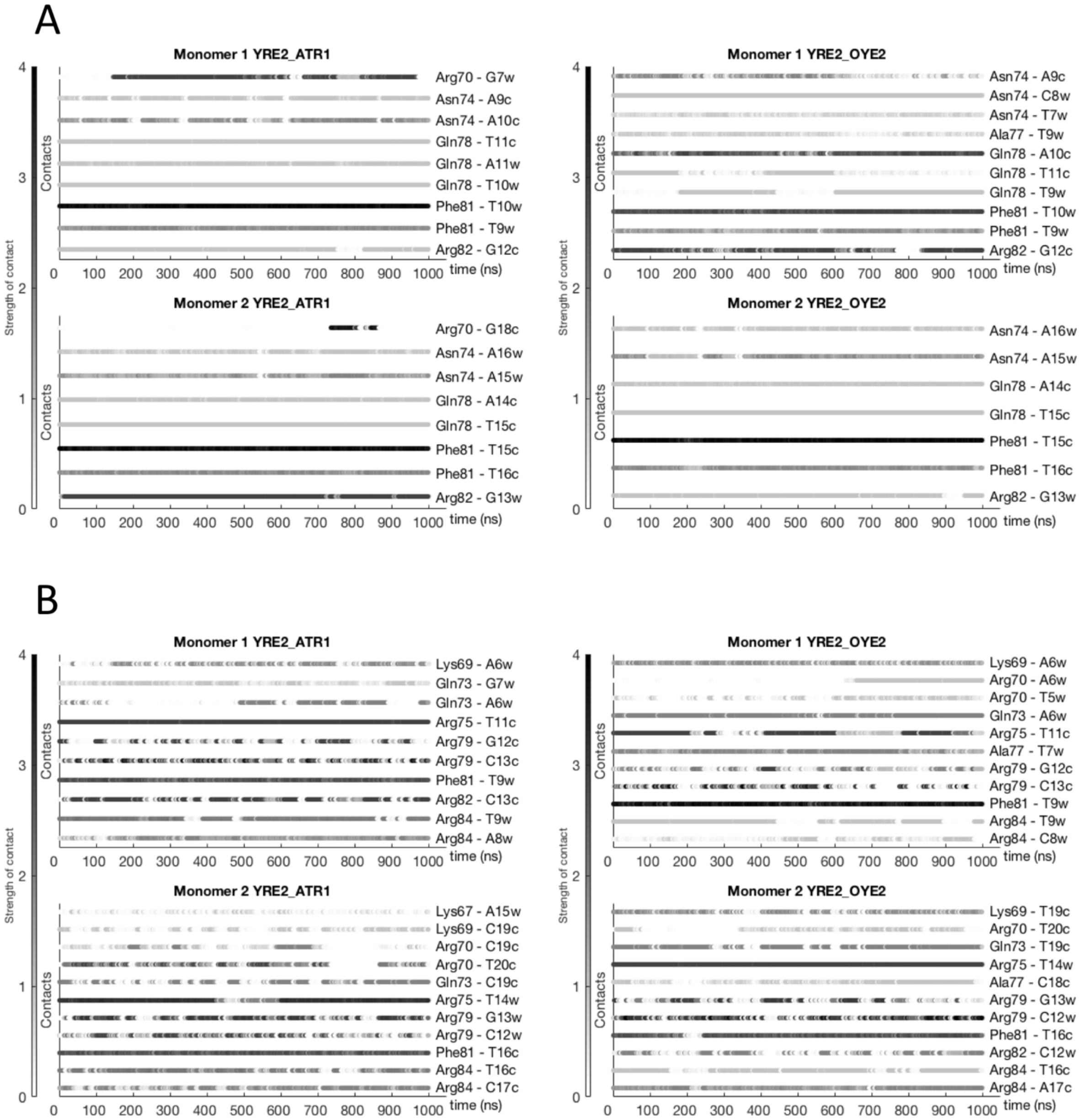


*Figure S3:* Dynamic interaction maps for Yap1 in complex with YRE2 in the ATR1- and OYE2-environments. The maps illustrate the strenght of **A.** specific and **B.** nonspecific Yap1-DNA contacts as a function of time. Indices “w” and “c” reffer to Watson- and Crick-DNA strands.

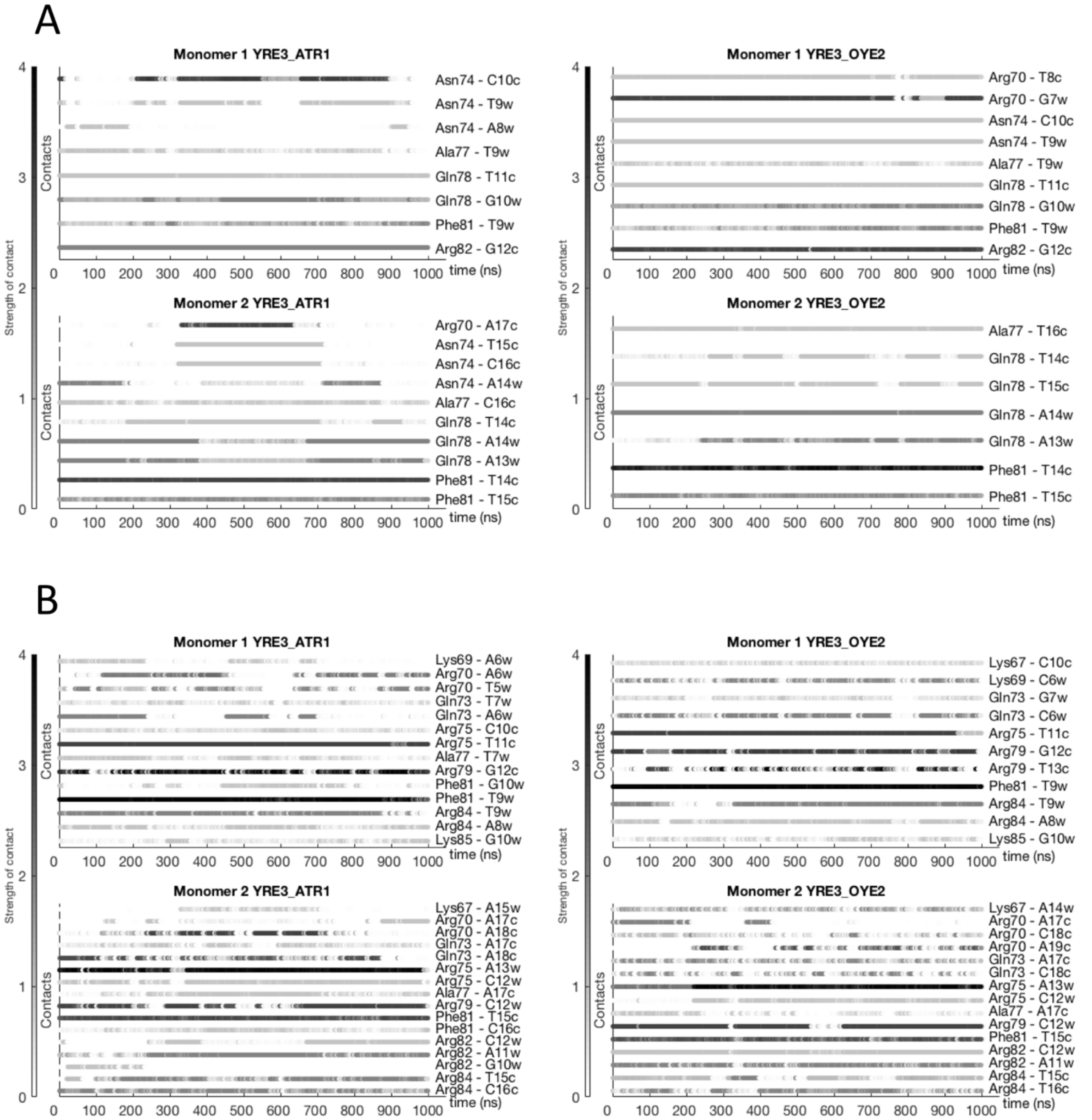


*Figure S4:* Dynamic interaction maps for Yap1 in complex with YRE3 in the ATR1- and OYE2-environments. The maps illustrate the strenght of **A.** specific and **B.** nonspecific Yap1-DNA contacts as a function of time. Indices “w” and “c” reffer to Watson- and Crick-DNA strands.

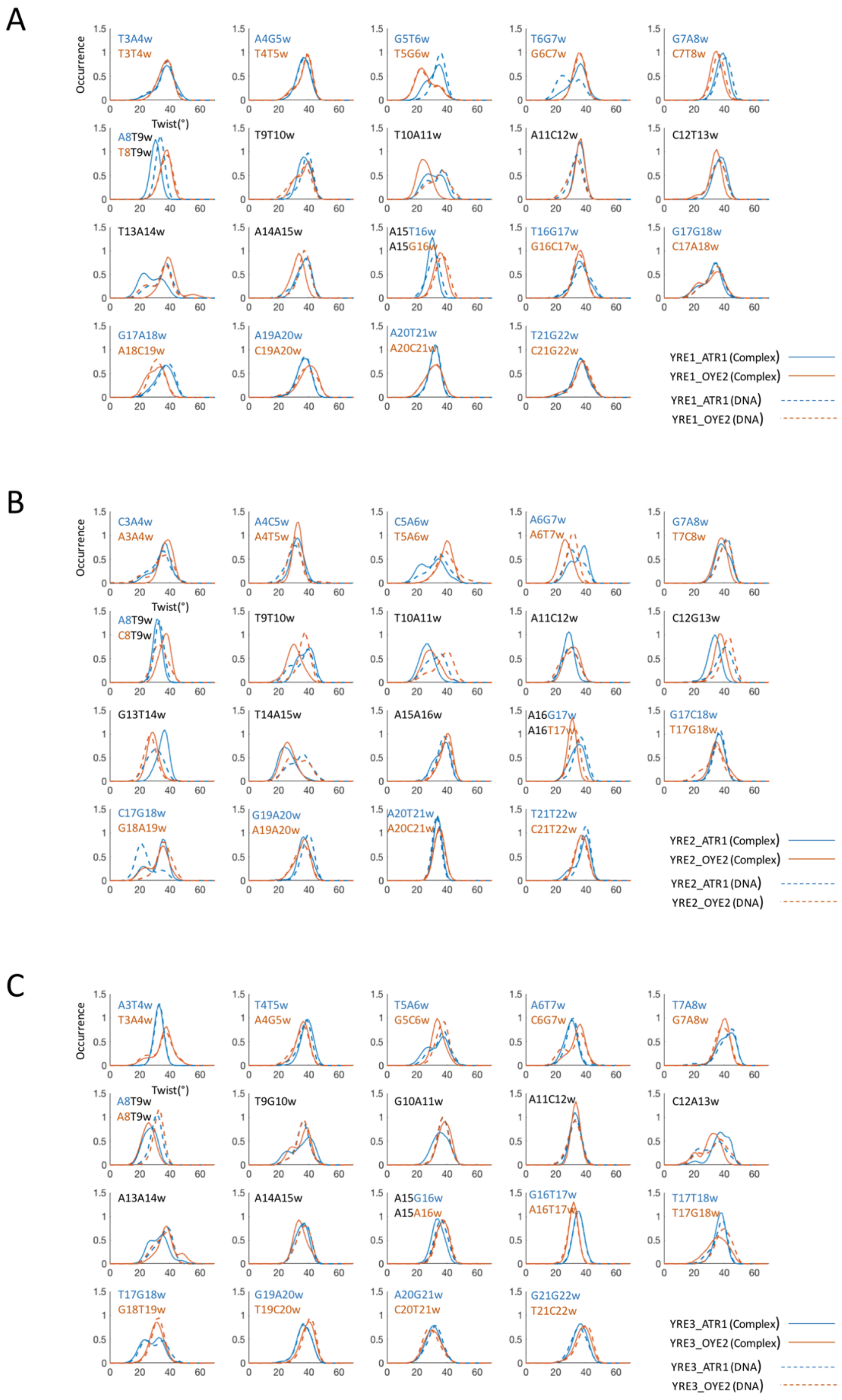

*Figure S5:* Normalised twist distributions for free DNA and DNA in complex with Yap1 for the three studied YREs **A.** Yap1-YRE1: TTACTAA, **B.** Yap1-YRE2: TTACGTAA, **C.** Yap1-YRE3: TGACAAA, in two genomic environments

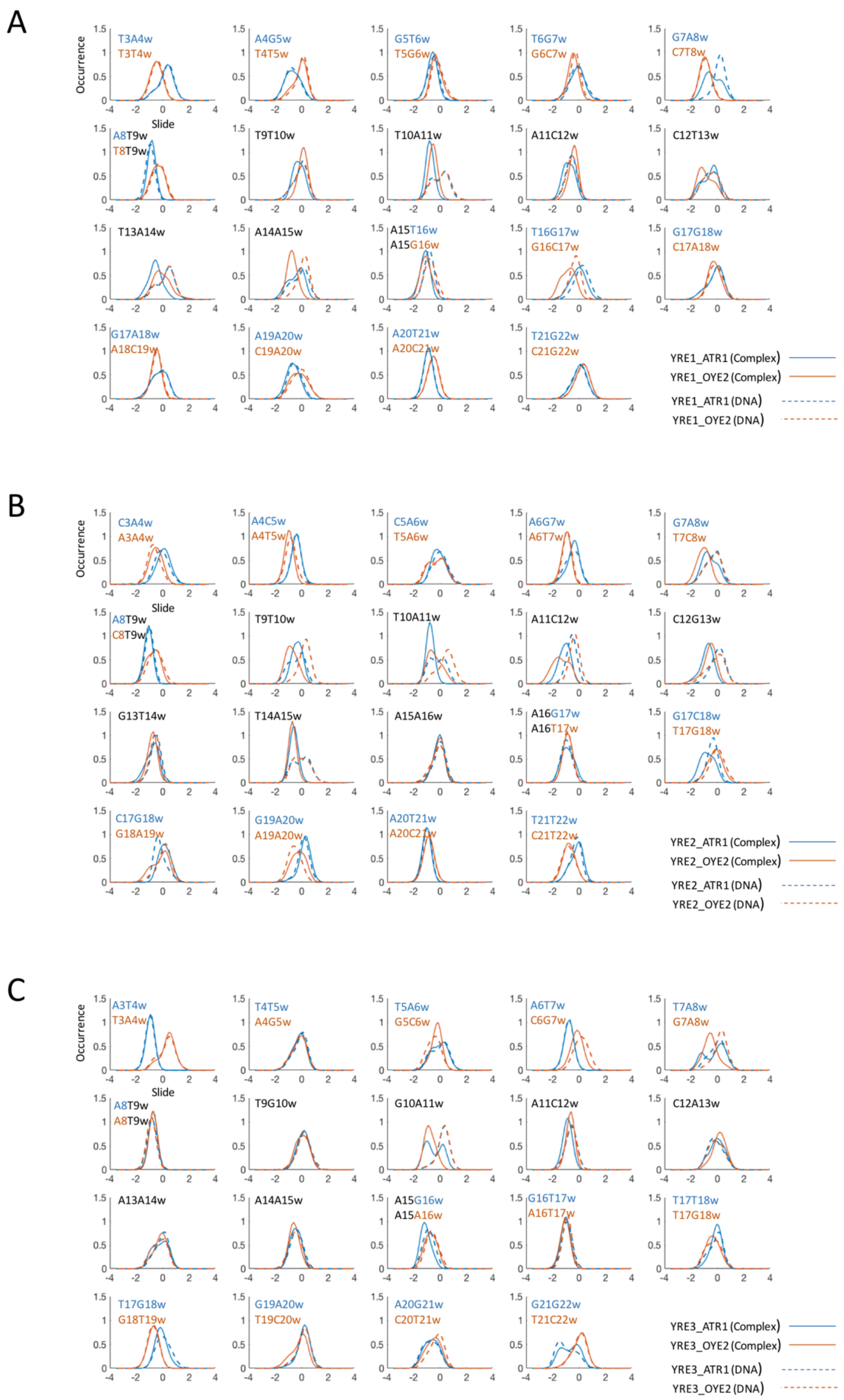


*Figure S6:* Normalised slide distributions for free DNA and DNA in complex with Yap1 for the three studied YREs **A.** Yap1-YRE1: TTACTAA, **B.** Yap1-YRE2: TTACGTAA, **C.** Yap1-YRE3: TGACAAA, in two genomic environments

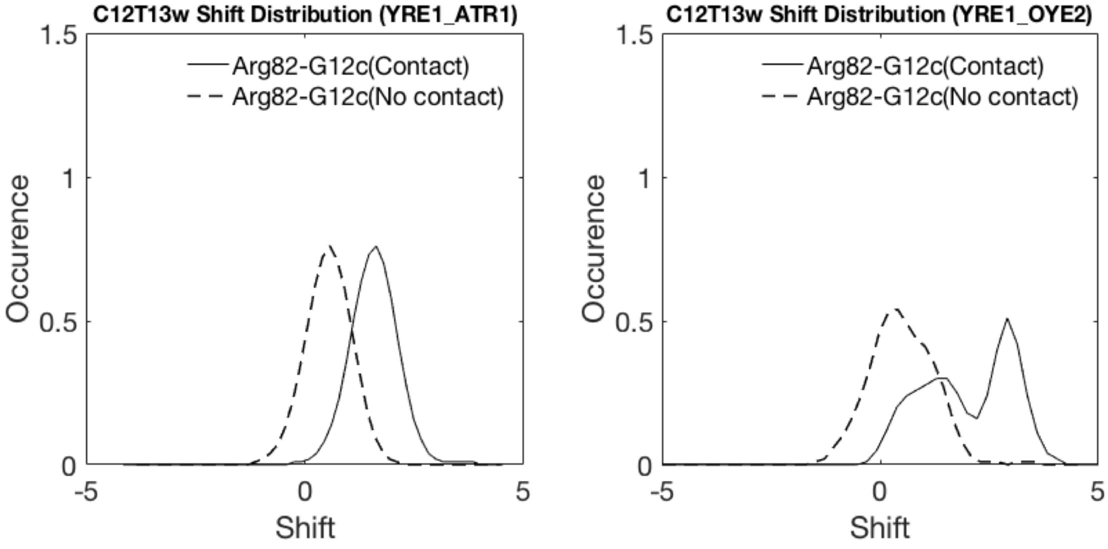


"w"-5′- TGCT / TGGA – TTACTAA – GCAC / ACCT -3′

5 6 7 8 9 10 11 12 13 14 15 16 17 18 19

"c"-3′- CACT / ACGA – AATGATT – GTGA / CGTG -5′

*Figure S7:* Shift distributions for the YRE1 b.p. step, C12T13w (TTA**CT**AA) in precence (black lines) and not precense (dashed lines) of the Arg82-G12c specific contact. The YRE1 DNA sequence is shown with the four adjacent flanking nucleotides for the two genomic environments, ATR1 (blue) and OYE2 (orange). Indices "w" and "c" reffer to Watson- and Crick-DNA strands.

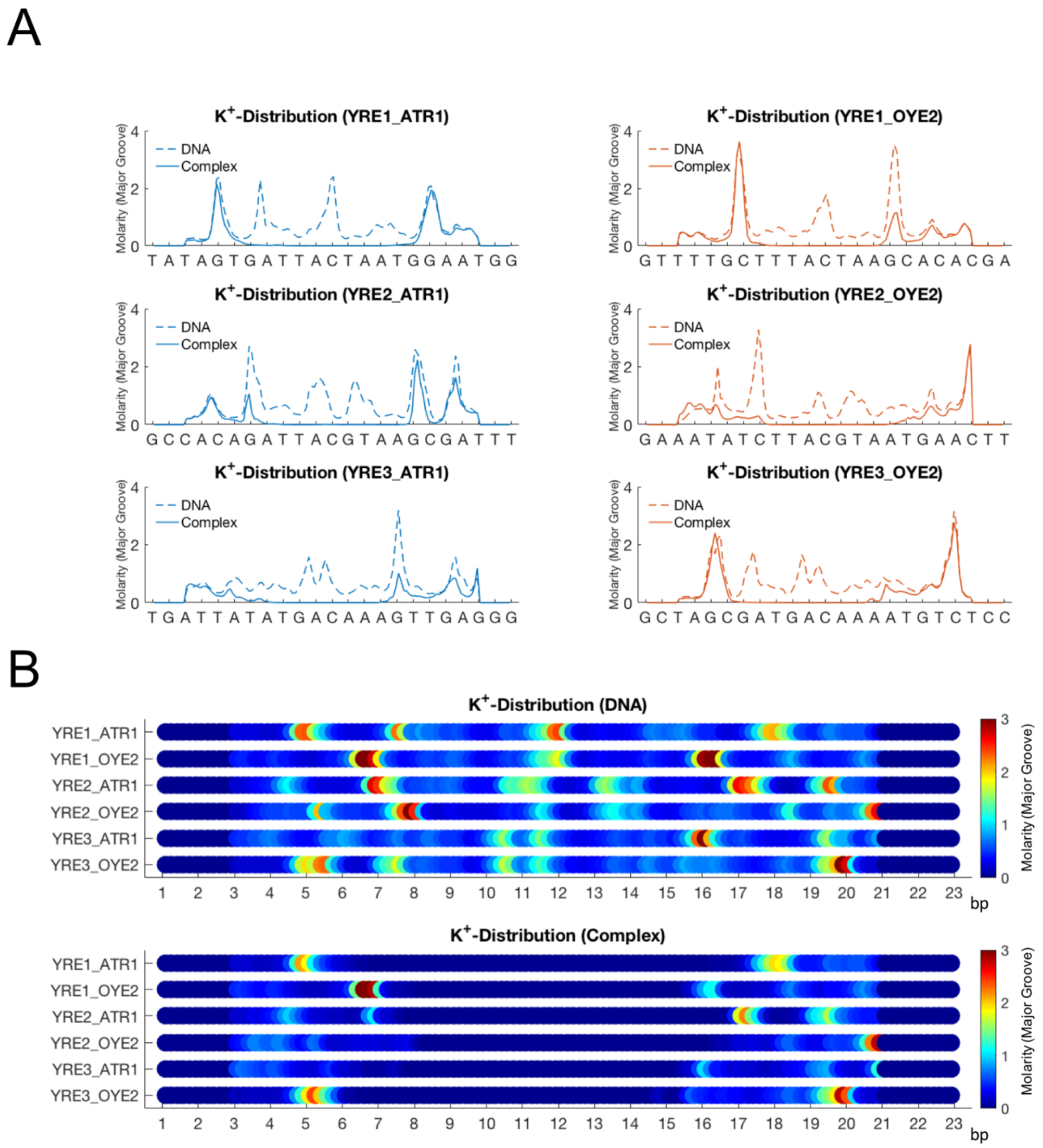


*Figure S8:* K+-distributions within DNA major groove along the DNA helix. **A.** Comparison of Yap1-bound DNA (thick lines) with unbound DNA (dashed lines) for the three YREs in the two genomic environments (ATR1: blue and OYE2: orange). **B.** Colormaps for the K+-distributions.

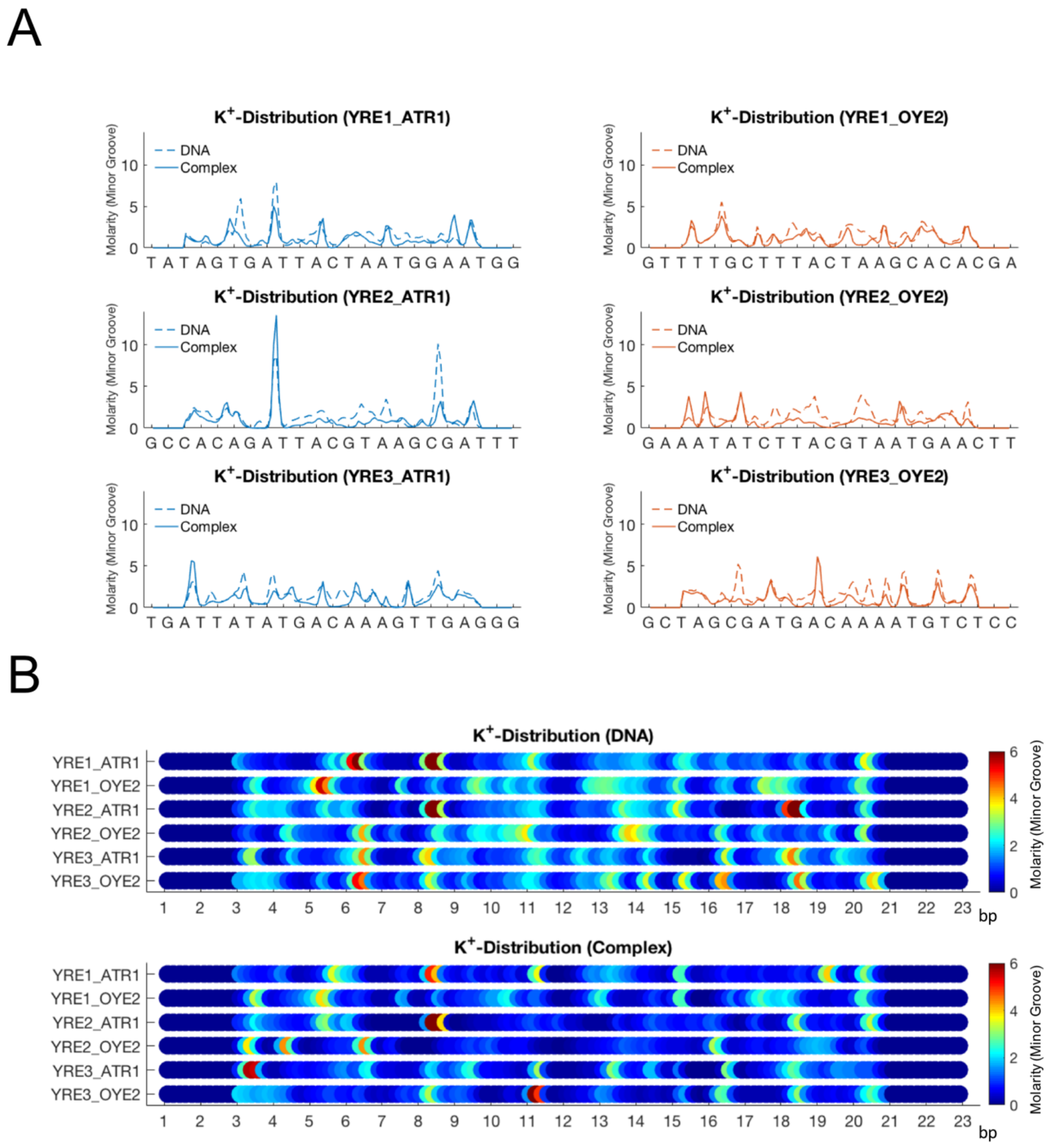


*Figure S9:* K+-distribution within DNA minor groove along the DNA helix. **A.** Comparison of Yap1-bound DNA (thick lines) with unbound DNA (dashed lines) for the three YREs in the two genomic environments (ATR1: blue and OYE2: orange). **B.** Colormaps for the K+-distributions.

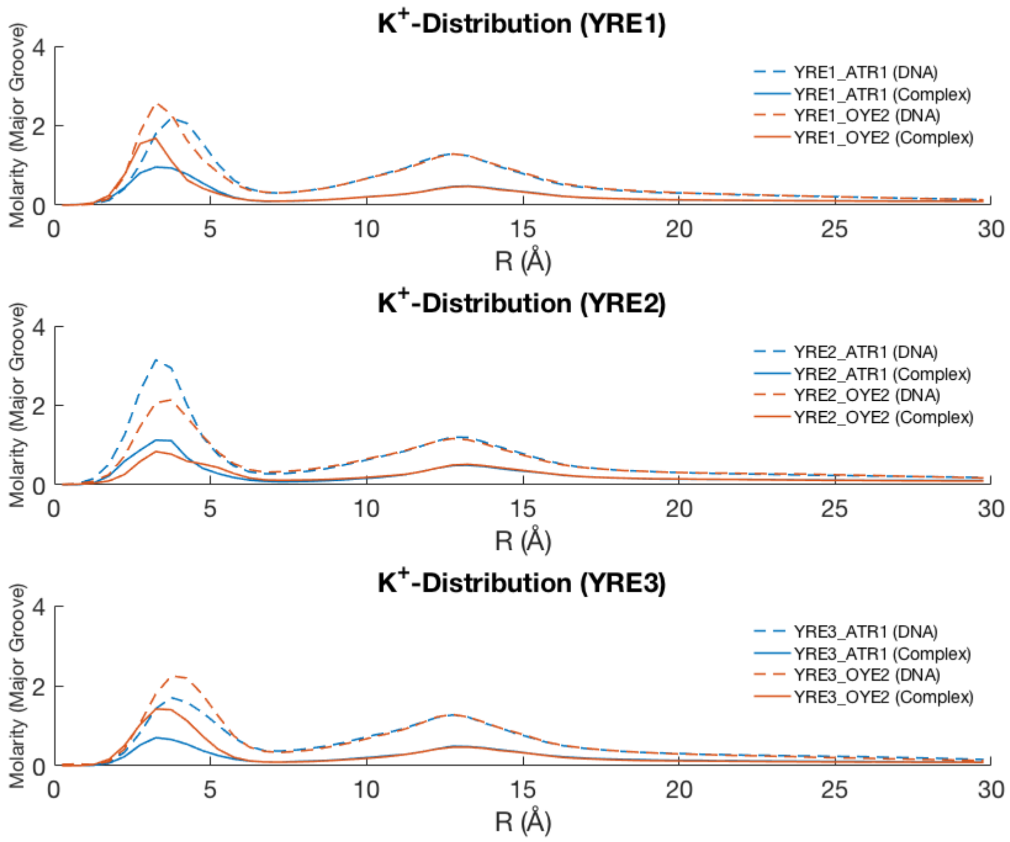


*Figure S10:* K+-distribution for DNA major groove at distance R (Å) from the helical axis for the three YREs (Yap1-bound DNA: thick lines, and unbound DNA: dashed lines) in the two genomic environments (ATR1: blue and OYE2: orange). A distance R<10.25 Å (phosphorous radius) constitutes the K+-molarity for the internal region, that is within the groove.

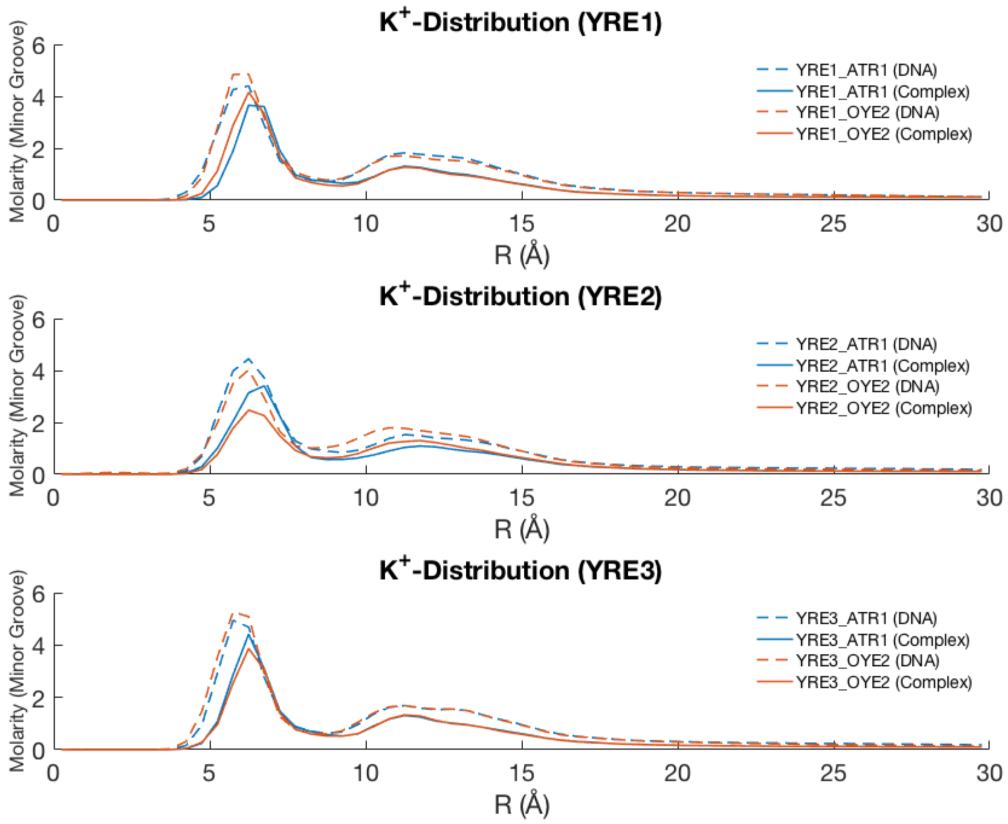


*Figure S11:* K+-distribution for DNA minor groove at distance R (Å) from the helical axis for the three YREs (Yap1-bound DNA: thick lines, and unbound DNA: dashed lines) in the two genomic environments (ATR1: blue and OYE2: orange). A distance R<10.25 Å (phosphorous radius) constitutes the K+-molarity for the internal region, that is within the groove.

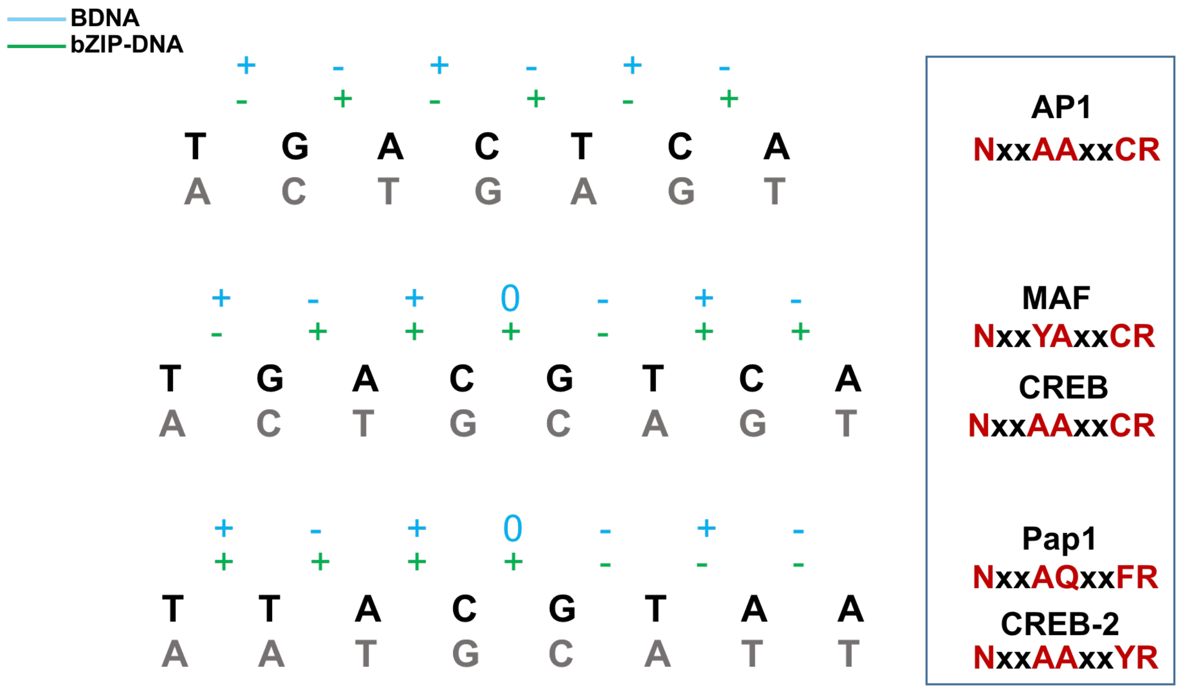


*Figure S12:* The sign of helical shift for b.p. steps within the three response elements for bZIP-bound DNA (derived from crystal structures) and B-DNA (derived from modelling tool JUMNA). The box to the right shows BZIP families (with the five-residue motif highlighted) that recognise the different response elements.
